# An improved *Nicotiana benthamiana* bioproduction chassis provides novel insights into nicotine biosynthesis

**DOI:** 10.1101/2023.03.06.531326

**Authors:** K Vollheyde, QM Dudley, T Yang, MT Oz, D Mancinotti, M Olivera Fedi, D Heavens, G Linsmith, M Chhetry, MA Smedley, WA Harwood, D Swarbreck, F Geu-Flores, NJ Patron

## Abstract

The model plant *Nicotiana benthamiana* is an increasingly attractive organism for the production of high-value, biologically active molecules. However, *N. benthamiana* accumulates high levels of pyridine alkaloids, in particular nicotine, which complicates the downstream purification processes. Here, we report the assembly of an improved *N. benthamiana* genome as well as the generation of low-nicotine lines by CRISPR/Cas9-based inactivation of berberine bridge enzyme-like proteins (BBLs). Triple as well as quintuple mutants accumulated 3-4 times less nicotine than the respective control lines. The availability of lines without functional BBLs allowed us to probe their catalytic role in nicotine biosynthesis, which has remained obscure. Notably, chiral analysis revealed that the enantiomeric purity of nicotine was fully lost in the quintuple mutants. In addition, precursor feeding experiments showed that these mutants cannot facilitate the specific loss of C6 hydrogen that characterizes natural nicotine biosynthesis. Our work delivers an improved *N. benthamiana* chassis for bioproduction and opens the possibility that BBLs are the sought-after coupling enzymes in nicotine biosynthesis.

## Introduction

*Nicotiana benthamiana* is a widely-used model plant native to Australia (Goodin *et al*., 2008; Bally *et al*., 2018). Following early observations of hypersusceptibility to viruses, *N. benthamiana* became valued for studying plant-pathogen interactions, and seeds were shared among the global plant science community (Bally *et al*., 2015). The hyper-susceptibility trait was eventually linked to a disruptive mutation in *RNA-dependent RNA polymerase 1* (*Rdr1*) present in ecotypes from arid regions (Bally *et al*., 2015). The mutation trades viral defense in favor of early vigor and thus confers an advantage in climates with low and unpredictable rainfall, where lifecycles must be completed in limited time (Bally *et al*., 2015; Karasov *et al*., 2017). Young *N. benthamiana* plants are also highly susceptible to *Agrobacterium tumefaciens*, a trait which has been exploited for transient gene expression in leaf tissues using a technique known as agroinfiltration (Chen *et al*., 2013b). In the last decade, facilities for the agroinfiltration of large numbers of plants have been constructed, thus enabling commercial-scale protein production (D’Aoust *et al*., 2010; Chen *et al*., 2013b; Lomonossoff & D’Aoust, 2016). Notably, *N. benthamiana* leaves can be agroinfiltrated using a complex mixture of *Agrobacterium* strains, leading to the transient co-expression of a multiplicity of genes. This has been exploited by the plant natural product community, who adopted it as the golden standard for enzyme discovery in the context of pathway elucidation (Geu-Flores *et al*., 2009; Gnanasekaran *et al*., 2015; Lau & Sattely, 2015; Reed *et al*., 2017; Stephenson *et al*., 2020; Davis *et al*., 2020; Dudley *et al*., 2022a). As more and more natural product pathways are elucidated this way, the facilities constructed for industrial protein production are likely to be repurposed for the production of high-value plant natural products such as terpenoids (Forman *et al*., 2022) or alkaloids (Nett & Sattely, 2021).

While *N. benthamiana* is a promising chassis for biomanufacturing, the yield and purity of protein products can be hampered by unintended proteolysis (Grosse-Holz *et al*., 2018). Similarly, small molecules can be further metabolized by oxidoreductases, hydrolases, and transferases that participate in *N. benthamiana’s* own metabolism or xenobiotic detoxification systems (van Herpen *et al*., 2010; Liu *et al*., 2011; Brückner & Tissier, 2013; Miettinen *et al*., 2014; Dong *et al*., 2016; Dudley *et al*., 2022a). In addition, like other *Nicotiana* species, *N. benthamiana* accumulates high amounts of nicotine and related pyridine alkaloids. Nicotine is the most abundant of these, with levels varying from 2.9 to 14.28 mg/g dry leaf weight (DW) (Sisson & Severson). This is of concern when producing therapeutic or dietary proteins, as nicotine is highly neuroactive and also potentially toxic in large amounts. In the production of thaumatin II, a small protein from the katemfe fruit used as sweetener, a chromatographic purification step was able to lower the levels of pyrimidine alkaloids down to levels encountered naturally in other Solanaceae vegetables such as tomato or pepper (*GRAS Notice 910*, 2020). We predict that the presence of nicotine will be more problematic for the industrial production of small molecules, in particular, other alkaloids with similar physio-chemical properties.

Nicotine biosynthesis occurs primarily in roots from where it is transported to aerial tissues via the xylem (Zenkner *et al*., 2019). On top of a basal level of production, nicotine biosynthesis can be induced by herbivore attack or by methyl jasmonate (MeJa) treatment (Steppuhn *et al*., 2004). The nicotine molecule is composed of two nitrogen-containing rings coupled directly to each other: a pyridine ring likely derived from nicotinic acid and a pyrrolidine ring evolutionarily derived from polyamine biosynthesis (Xu *et al*., 2017) (Figure 1). The pyridine branch is well characterized up to nicotinic acid mononucleotide, from which nicotinic acid is thought to derive (Shoji & Hashimoto, 2011). The pyrrolidine branch is fully characterized and gives rise to the *N*-methylpyrrolidinium cation, which is proposed to be coupled directly to a reduced & decarboxylated form of nicotinic acid (Shoji & Hashimoto, 2011). Finally, the coupled intermediate likely undergoes oxidation to give the final nicotine molecule (Figure 1). The details of these last steps of nicotine biosynthesis remain undefined; however, one important clue is that the hydrogen at position C6 of the pyrimidine ring in nicotinic acid is lost during biosynthesis (Dawson *et al*., 1960; Leete & Liu, 1973; Leete, 1978) (Figure 1). The related pyridine alkaloids anatabine and anabasine are also composed of two coupled nitrogen rings each; however the composition of the non-pyridine rings varies. In the case of anatabine, both rings are derived from the pyridine branch, while in the case of anabasine, one ring comes from the pyridine branch and the second one comes from a piperidine branch originating from the amino acid L-lysine (Figure 1). All three pyridine alkaloids possess a stereocenter at position C2 of the non-pyridine ring. In *N. tabacum*, the (*S*) forms predominate (Armstrong *et al*., 1998, 1999), most notably for nicotine, where the enantiomeric ratio reaches ca. 99.8% (Armstrong *et al*., 1999). In animals, the (*S*) enantiomer of nicotine is more toxic and displays greater pharmacological effects (Yildiz, 2004). The biosynthetic step that determines the (*S*) configuration remains to be determined, but it is conceivable that the coupling step is under stereoselective enzymatic control, thus producing the right stereochemistry that is retained during the final oxidation step.

**Figure 1.**
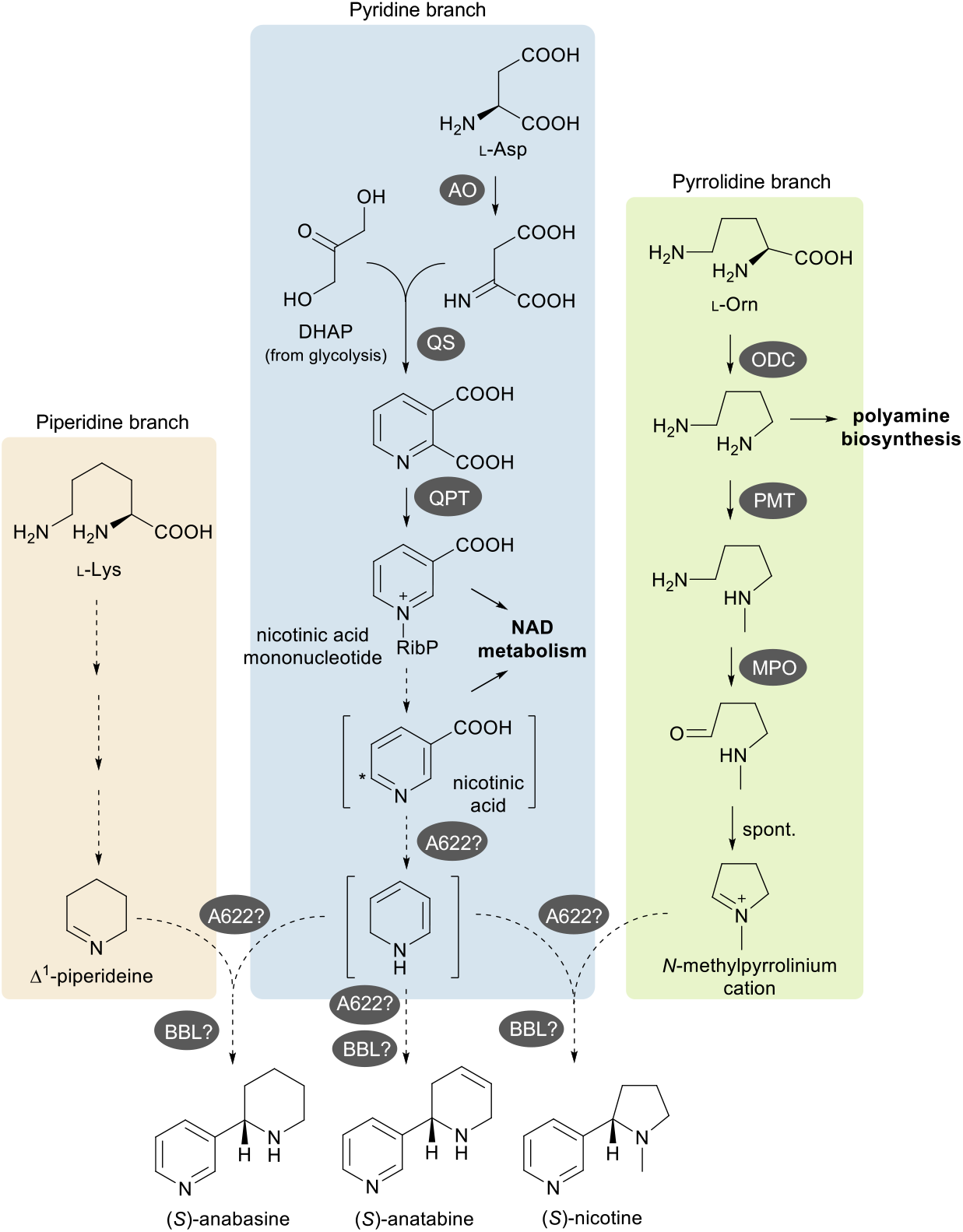
Schematic diagram of pyridine alkaloid biosynthesis in *Nicotiana* spp. The three colored boxes highlight the different biosynthetic branches involved. Full arrows represent single characterized steps, and dotted arrows represent one or more putative steps. The intermediates shown in brackets are putative. The asterisk in nicotinic acid indicates position C6, whose hydrogen is lost during biosynthesis (Dawson *et al*., 1960). Enzymes are represented by dark gray ovals. For the piperidine branch (left), question marks on enzyme names indicate that the enzymes have not yet been cloned. For A622 and BBL, the question marks indicate uncertainty about precisely which reactions these enzymes catalyze. AO: aspartate oxidase; QS: quinolinate synthase; QPT quinolinate phosphorybosyltransferase; ODC: ornithine decarboxylase; PMT: putrescine methyltransferase; MPO: methylputrescine oxidase; A622: PIP-family reductase; BBL: berberine bridge enzyme-like protein; L-Lys: L-lysine; L-Orn: L-ornithine; RibP: ribose phosphate residue; NAD: nicotinamide adenine dinucleotide.

Two enigmatic enzymes have been implicated in the late biosynthetic steps mentioned above for all pyridine alkaloids. The first one is a reductase of the phosphatidylinositol phosphate (PIP) family called A622. Conditional suppression of two *A622* copies in *N. tabacum* hairy roots via RNAi resulted in cell growth inhibition, lower levels of pyridine alkaloids, and accumulation of a nicotinic acid *N-*glucoside as well as the *N*-methylpyrrolidinium cation (Kajikawa *et al*., 2009). This suggests that A622 catalyzes the putative reduction & decarboxylation of nicotinic acid and/or the coupling step (Kajikawa *et al*., 2009). However, there are no reports showing the catalytic properties of A622. The second implicated enzyme is a berberine bridge enzyme-like protein (BBL). Simultaneous suppression of several *BBL* copies in *N. tabacum* plants and hairy roots via RNAi led to a reduction in pyridine alkaloid levels and accumulation of dihydrometanicotine (DMN) (Kajikawa *et al*., 2011) (Lewis *et al*., 2015). DMN itself does not appear to be a substrate of BBLs, as recombinant proteins expressed in *Pichia pastoris* exhibited no activity towards it (Kajikawa *et al*., 2011). Further evidence for BBL involvement has come from the simultaneous inactivation of two or three *N. tabacum BBL* copies via chemical mutagenesis, which reduced the nicotine content up to 13-fold (Lewis *et al*., 2015). Later, CRISPR/Cas9 technology was used to introduce mutations into all six *BBL* copies found in the *N. tabacum* genome, producing lines reported to be nicotine free (Schachtsiek & Stehle, 2019). By contrast, a follow-up study on the chemically mutagenized double and triple *BBL* mutants showed that further inactivation of the remaining *BBL* genes by CRISPR/Cas9 technology did not result in any further nicotine reductions (Lewis *et al*., 2020).

While the involvement of A622 and BBL in nicotine biosynthesis is undisputed, the exact reactions that they catalyze are yet to be determined. In this study, we report an improved assembly of the *N. benthamiana* genome and identify five *BBL* genes. We use CRISPR/Cas9 technology to generate low-nicotine lines with loss-of-function mutations in individual and multiple combinations of *BBL* genes. Surprisingly, analysis of lines with no functional BBL revealed that the residual nicotine was fully racemic. Furthermore, precursor feeding studies indicated that the mutants did not undergo the specific loss of C6 hydrogen as observed in the control lines. Our work provides an improved low-nicotine *N. benthamiana* chassis for bioproduction and sheds light into the precise catalytic role of BBL enzymes in nicotine biosynthesis.

## METHODS

### Plant growth

*N. benthamiana* plants were grown in a controlled environment room with 16 hr light, 8 hr dark at 22 °C, 80% humidity, and ~200 μmol/m^2^/s light intensity.

### Production of paired end reads

DNA was purified from young leaf tissue using a modified CTAB-based protocol (Michael *et al*., 2018). Briefly, 5 g of frozen powdered leaf tissue was incubated with 20 mL CTAB lysis buffer (100 mM Tris-HCl, 2% CTAB, 1.4 M NaCl, 20 mM EDTA, pH 8.0) containing 20 μg/mL proteinase K for 20 minutes at 55 °C. Aliquots of 2.5 mL were extracted with 0.5 volumes of chloroform followed by an equal volume of 25:24:1 phenol:chloroform:isoamyl alcohol (PCI). DNA was precipitated from the upper aqueous phase with 0.7 volumes of ice-cold isopropanol. The pellet was washed with ice-cold 70% ethanol and air-dried at room temperature before resuspension in TE (10 mM Tris-HCl pH 7.5, 1 mM EDTA) containing 1.4 mg/mL RNase A. RNase A was then removed by two extractions with PCI and DNA was precipitated as above and resuspended in TE. Samples were pooled on a Genomic DNA Clean & Concentrator-10 column (Zymo Research, Irvine, CA, USA) and eluted in 50 μL DNA elution buffer (10 mM Tris-HCl, pH 8.5, 0.1 mM EDTA). Amplification free, Illumina compatible libraries were constructed using the Hyper Prep kit (Kapa Biosystems, Wilmington, MA, USA). DNA was sheared to 1 Kbp using an S2 column (Covaris, Woburn, MA, USA) and Kapa Pure Beads (Kapa Biosystems) were used to remove molecules of less than <500 bp. DNA molecules were end repaired, A-tailed and appropriate Illumina-compatible indexed adapters were ligated. Library QC was performed using a BioAnalyzer high sensitivity chip (Agilent, Santa Clara, CA, USA) and the concentration of viable library molecules measured using qPCR. For sequencing, libraries were loaded at 9 pM based on the qPCR concentration with an average molecule size of 625 bp. Libraries were sequenced on an Illumina HiSeq 2500 yielding 272,102,680 250 bp reads. Paired end reads were deposited in the European Nucleotide Archive (ENA) under project number PRJEB37024 (accession numbers: ERR3971933 and ERR3971934).

### Production of Oxford Nanopore Technologies (ONT) long-reads

DNA for sequencing on the MinION (Oxford Nanopore, Oxford, UK) was purified from leaf tissue sampled from an individual plant, using a modified version of the CTAB protocol above. Briefly 1.0 g of frozen powdered leaf tissue was mixed with 3 mL CTAB lysis buffer containing 20 μg/mL proteinase K and incubated at 55 °C at 500 rpm. After 15 minutes, 20 μL of 100 mg/mL RNaseA was added and incubation continued for 15 minutes before a further 20 μL of 100 mg/mL RNaseA was added. Extractions with chloroform and PCI were performed as above except that wide-bore pipette tips were used at all stages. DNA was precipitated and resuspended in 25 μL TE. Yields (8-21 μg at 350-900 ng/μL) were determined using the Qubit fluorometer (ThermoFisher, Waltham, MA, USA). Fragments of >40 Kbp were collected for library preparation using BluePippin pulsed-field electrophoresis (Sage Science, Beverly, MA, USA). DNA was stored at 4 °C overnight before proceeding directly to library preparation. Libraries were constructed and sequenced using the Oxford Nanopore Technologies (ONT) SQK-LSK109 library construction kit and FLO-106 flow cells according to the manufacturers’ instructions. A total of nine MinION flow cells were used yielding between 3 and 20 Gbp per flow cell. Two flow cells were loaded with libraries constructed from non-fragmented DNA. Two flow cells were loaded with libraries constructed from DNA sheared to 40 Kbp using the Hydroshear (Digilab, Hopkinton, Massachusetts, USA). Molecules <25 Kbp were removed using a Blue Pippin (Sage Scientific, Massachusetts, USA) with a 0.75% cassette. Two flow cells were loaded with libraries constructed from DNA sheared to 40 Kbp and molecules of <15 Kbp removed. Three flow cells were loaded with libraries constructed from DNA sheared to 20 Kbp using a G-tube (Covaris, Woburn, Massachusetts, USA). Reads were deposited in the ENA under project number PRJEB37025, accession numbers ERR3971506-509 and ERR3972081-83.

### Production of Chromium linked reads

DNA for Chromium linked-read sequencing (10X Genomics) was purified from leaf tissue sampled from an individual plant using the GE Healthcare Nucleon™ PhytoPure™ Genomic DNA Extraction Kit (Fisher Scientific, Loughborough, Leicestershire, UK) following the manufacturer’s instructions except that, following precipitation with an equal volume of ice-cold isopropanol, DNA was collected by centrifugation at 1,300 x g for 10 min at 4 °C and the resulting DNA pellet washed with 1 mL 70% ethanol air-dried and resuspended in 40 μL Low TE (10 mM Tris-HCl pH 7.5, 0.1 mM EDTA). Purified DNA was quantified by Qubit (21 ng/μL) and analyzed on the Femto Pulse System (FP-1002-0275, Agilent, Santa Clara, CA, USA) with the dominant peak at 150,000 bp and 67% of the material being greater than 50 Kbp. The Chromium Controller instrument (10x Genomics, Pleasanton, California, USA) was used to produce a barcoded linked read library using 1.25 ng DNA input and the Chromium™ Genome Library Kit & Gel bead Kit v2 (120258, 10x Genomics) following the Chromium Genome Reagent Kits Version 2 User Guide (CG00043, 10x Genomics). Library yield was quantified using the Qubit dsDNA HS assay and insert size was determined using the 2100 Bioanalyzer High Sensitivity DNA chip (5067-4627, Agilent Technologies, Santa Clara, California, USA) and verified by qPCR. The library was diluted to 0.5 nM with EB in 18 μL and spiked with 1% PhiX Control v3 (Illumina) before being prepared for loading using a NovaSeq XP 2 lane kit v1.5 (20043130, Illumina). This was sequenced with 150 paired-end reads on an Illumina NovaSeq 6000 with NVCS 1.7.5 and RTA v3.4.4 on one lane of a NovaSeq S4 v1.5 flow cell with accompanying reagent cartridges (20028312, Illumina) yielding 891 Gb (2,971,099,042 reads). Sequencing data was demultiplexed and converted from base call (BCL) files to FASTQ files using the Illumina bcl2fastq2 conversion software (bcl2fastq version. 2.20.0), allowing for a one base-pair mismatch to the index sequence. Reads were deposited in the ENA under project numbers PRJEB37026 (accession number ERR3972084 and ERR3972085) and ERX10379414.

### Genome assembly

A combination of Chromium linked-reads and ONT long-reads were utilized to generate the *N. benthamiana* genome assembly. Chromium linked-reads were assembled using Supernova v2.1.1 (https://bio.tools/supernova) (Weisenfeld *et al*., 2017) with the reads subsampled to 56x raw coverage (-- maxreads= 1136580012) as recommended by the Supernova documentation, resulting in an initial draft assembly with a N50 of 5.4 Mb. The assembly was scaffolded using ONT long-reads. To improve the quality of the scaffolding process, ONT long-reads were polished using the Illumina paired-end short reads with the tool ratatosk (v0.7.6.3; https://github.com/DecodeGenetics/Ratatosk) (Holley *et al*., 2021) with default options. Long read scaffolding was performed with ntLink (v1.3.4; https://github.com/bcgsc/ntLink) in gap-filling mode, with a k-mer size of 40 for mapping (--k=40), and a requirement that at least four reads validate the joined contigs (--a=4). This process was repeated five times. Fragments of assembled chloroplast and mitochondrial genome were identified and excluded from the final assembly. Specifically, Minimap2 v2.22 (Li, 2018) was used to align a reference chloroplast genome (accession number: MF577082.1) and multiple mitochondrial genomes of related *Nicotiana* species (accession numbers: MN651321.1, MN651322.1, and MN651323.1) to the scaffolded assembly. Contigs with greater than 99% identity and with comparable size to the reference chloroplast genome were removed, while all contigs that mapped to the three mitochondrial genomes and that had more than 70% of their sequence covered were considered mitochondrial content and also removed from the assembly.

To further identify and remove any remaining contamination, the FCSx contamination pipeline (v0.3.0; https://github.com/ncbi/fcs) was employed, which identified several contigs of bacterial and primate origin. These contigs were subsequently removed from the final assembly. To assess the completeness of the final assembly, (Benchmarking Universal Single-Copy Orthologs) BUSCO v5.3.2 (https://gitlab.com/ezlab/busco/-/releases#5.4.4) (Simão *et al*., 2015) was run with metaEuk as the aligner against the eudicots_odb10 database. Additionally, a Merqury v1.3 (Rhie *et al*., 2020) analysis was performed to obtain the k-mer completeness and the QV of the assembly, which provides a complementary assessment metric to the BUSCO score. The final scaffolded assembly can be accessed from the following url: https://opendata.earlham.ac.uk/opendata/data/Patron_2023-03-01_Nicotiana_benthamiana_genome_assembly/.

### Expression analysis

For expression analysis, wild type plants were treated with a foliar spray of 2.5 mM MeJa in dimethyl sulfoxide (DMSO) or DMSO (control plants) seven days before sampling. Total RNA was isolated from 100 mg of fresh root tissue sampled from three individual plants (biological replicates) using the Spectrum Plant RNA Purification Kit (Merck). RNA was treated with RNase-free DNase Set (Qiagen) and cDNA was synthesized from 300 ng of total RNA using the M-MLV Reverse Transcriptase (ThermoFisher Scientific). SYBR^®^ Green JumpStart™ Taq ReadyMix™ (Merck) was employed for reporting successful amplification using a QuantStudio 6 Pro real-time PCR system (Applied Biosystems, Waltham MA) with specific primer pairs for each target *NbBBL* gene. Primer sequences used for expression analysis are provided in Table S1. *NbEF1a* was amplified as an internal reference (Liu *et al*., 2012) for relative gene expression. Four technical replicates were performed for each sample and primer pair.

### Cas9-mediated targeted mutagenesis by Agrobacterium-mediated transformation

A binary vector for Cas9-mediated targeted mutagenesis via *Agrobacterium tumefaciens*-mediated transgenesis was assembled using the plant modular cloning toolkit (Engler *et al*., 2014) as previously described (Dudley *et al*., 2022b). Two single guide RNA (sgRNA) expression cassettes were assembled by amplifying the sgRNA scaffold with an extended stem (Chen *et al*., 2013a) from plasmid pEPOR1CB0022 (Addgene #117537) and integrating the spacer sequence for each target as a 5’ extension of the forward primer. Primer sequences used to amplify sgRNAs are provided in Table S2. The resulting amplicons were assembled in Level 1 acceptors with an AtU626 promoter (pICSL90002, AtU6-26 Addgene#68261). The final construct was assembled by combining the two Level 1 sgRNA cassettes with cassettes for resistance to kanamycin pICSL11024 (Addgene#51144) and constitutive expression of SpCas9 (pEPQD1CB0001, Addgene #185625). All four Level 1 cassettes were assembled in a one-step reaction into the Level 2 acceptor plasmid (pAGM4723 Addgene #48015) as previously described (Dudley *et al*., 2022b). The efficacy of the resulting binary construct was tested by transient infiltration as previously described (Dudley *et al*., 2022b). The resulting constructs were transformed into *A. tumefaciens* strain AGL1 and used for transformation of *N. benthamiana* as previously described (Dudley *et al*., 2022a).

### Cas9-mediated targeted mutagenesis using a TRV2 viral vector

Spacer sequences for selected targets were incorporated into pEPQD0KN0750 (Addgene #185627) by Golden Gate assembly into AarI sites as previously described (Dudley *et al*., 2022a). Primer sequences are provided in Table S3. Constructs were transformed into *A. tumefaciens* GV3101 and infiltrated into transgenic *N. benthamiana* plants constitutively expressing Cas9 (Cas9 Benthe 193.22 T5 Homozygous; gratefully received from Dan Voytas) together with a strain containing pTRV1 (Addgene #148968) as previously described (Ellison *et al*., 2020; Dudley *et al*., 2022a). Plants were allowed to grow for 13 weeks before samples of leaf tissue were taken from two different stems.

### Genotyping of gene-edited lines

Genomic DNA was isolated from the leaves of T_0_ plants generated by agrobacterium-mediated transformation or from leaves sampled from E_0_ plants infiltrated with TRV viral vectors as previously described (Dudley *et al*., 2022a). Target loci were amplified using a proof-reading polymerase (Q5^®^ High-Fidelity DNA Polymerase, New England Biolabs, Ipswich, MA; Phusion™ High-Fidelity DNA Polymerase, Thermo Fisher Scientific, Waltham, USA) and primers flanking the target sites. Primer sequences used to genotype edited lines are provided in Table S4. Amplicons were sequenced by Sanger sequencing (Eurofins, Luxembourg). Seeds were harvested from primary transgenic plants (T_0_) in which mutations were detected at one or more target loci. Seeds from plants infiltrated with TRV viral vectors (E_0_ plants) were harvested from seed pods on stems in which mutations had been detected. The resulting T_1_ and E_1_ plants were grown and the genotype of each target loci was confirmed. Stable, non-chimeric lines with homozygous or biallelic mutations at one or more target loci were selected for further analysis and T2/E2 seed collection. The genotypes of all *BBL* genes were confirmed in the T2/E2 generation.

### Leaf alkaloid analysis of gene-edited lines

T2/E2 plants were grown in soil in a greenhouse under long day conditions (16 h light) and day/night temperatures of 20/19 °C. After 34 days, six 1-cm leaf discs per plant representing three different leaves (two leaf discs per leaf) were harvested. The leaf discs were snap-frozen and stored at −70°C until further use (uninduced samples). Subsequently, alkaloid biosynthesis was induced by spraying leaves with 0.1% (*v/v*) MeJa solution containing 0.1% (*v/v*) Tween 20 (source of MeJA: Duchefa Biochemie, Haarlem, Netherlands). Five days after induction, another six leaf discs were harvested from the same leaves as before, snap-frozen and stored at −70°C. Frozen leaf discs from uninduced and induced leaves were subjected to metabolite extraction, LC-MS analysis, and, for three of the lines, (*R*)- and (*S*)-nicotine analysis as described below.

### Precursor feeding experiments

Seedlings were grown using a hydroponic system illustrated in Figure S1. For each sample, 20 seeds were germinated and grown on cotton gauze (28 thread) glued onto ¾-inch rubber O-rings fitted onto the wells of 12-well plates. Seedlings were grown under long day conditions (16 hr light) in tap water that was refilled when needed. Clear plate lids were placed on the plates (touching the O-rings) for the first five days. Alkaloid biosynthesis was induced 17 days after placing the seeds on the cotton gauze. For the induction, two filter papers with 5 μL MeJa (Duchefa Biochemie, Haarlem, Netherlands) each were placed next to the plates, and one large cover (clear plastic) was placed over all plates, facilitating a closed environment. Two days later, the tap water was replaced with either 0.8 mM D4-nicotinamide (D4 98%, Cambridge Isotope Laboratories, Tewksbury, USA) or 0.8 mM unlabeled nicotinamide, both solutions in tap water. The respective nicotinamide solutions were refilled two days after the start of the feeding (continuous feeding). Fresh MeJa was added to the filter papers at the start of the feeding and also during the refill. Seedlings were harvested five days after the commencement of feeding. All seedlings from one sample were pooled and rinsed three times with 1 mL ultrapure water. Rinsed seedlings were snap-frozen and stored at −70°C. Frozen samples were subjected to metabolite extraction and LC-MS analysis as described below.

### Root alkaloid analysis of gene-edited lines

To examine DMN formation in roots, seedlings were grown in a hydroponic system (Figure S1) as described above, except that 6-well plates with 1¼-inch O-rings were used instead, as well as 40 seeds per sample. 35 days after placing the seeds on the cotton gauze, alkaloid biosynthesis was induced using the MeJa filter paper method described above. Three days after the start of the induction, the wells were refilled with tap water and fresh MeJa was added to the filter papers. The seedlings were harvested five days after the start of the induction. For harvesting, seedlings were rinsed six times with 500 μL ultrapure water, and the roots were cut off, snap-frozen, and stored at −70°C. Frozen samples were subjected to metabolite extraction and LC-MS analysis as described below.

### Metabolite extraction

Frozen samples of leaf discs, whole seedlings, or seedling roots were lyophilized using a freeze dryer (CooleSafe™, Scanvac). Freeze-dried tissues were homogenized using chrome balls and a TissueLyzer (Qiagen). Metabolites were extracted with 60% (*v/v*) methanol containing 0.06% (*v*/*v*) formic acid and either 75 ppm caffeine (for leaf discs) or 25 ppm caffeine (for whole seedlings and seedling roots). The proportions of dry tissue (mg) to extractant (μL) varied depending on the tissue source: 4 mg/100 μL for leaf discs, 4 mg/300 μL for whole seedlings, and 4 mg/1000 μL for seedling roots. After 1 hr incubation at room temperature with constant shaking (1,200 rpm), samples were spun down for 1 min at 16,200 *x g*. Supernatants were diluted in ultrapure water at a proportion of 1:15 in the case of leaf discs or 1:5 in the case of whole seedlings and seedling roots. Diluted samples were filtered through a 0.22 μm PVDF filter (MultiScreen_HTS_ GV Filter Plate, 0.22 μm, clear, non-sterile, Merck Millipore, Billerica, USA) and stored at −70°C until further analysis.

### LC-MS analysis

Methanolic extracts were analyzed via reversed-phase LC-MS on a Dionex UltiMate 3000 Quaternary Rapid Separation UHPLC+ focussed system (ThermoFisher Scientific) coupled to an ESI QTOF Compact mass spectrometer (Bruker, Bremen, Germany). Compounds were separated on a Kinetex^®^ 1.7 μm EVO C18 100 Å column (100 x 2.1 mm, Phenomenex, Torrance, USA) applying an eluent flow rate of 0.3 mL/min and a column temperature of 40°C. Mobiles phases A and B consisted of 10 mM ammonia bicarbonate (pH 9.2) in water and acetonitrile, respectively, and the elution profile consisted of 0-1 min 2% B (constant), 1-16 min 2-25% B (linear), 16-24 min 25-65% B (linear), 24-26 min 65-100% B (linear), 26-27 min 100% B (constant), 27-27.5 min 100-2% B (linear), and 27.5-33 min 2% B (constant). Mass spectra were obtained in positive ionization mode with automatic MS/MS acquisition and using the following parameters: capillary voltage 4500 V, end plate off set 500 V, dry temperature 250°C, dry gas nitrogen flow rate 8.0 L/min, and nebulizing gas pressure 2.5 bar. MS spectra were recorded in an *m/z* range from 50-1000 Da (spectra rate: 6 Hz). Internal mass calibration was facilitated with Na-formate clusters. The injection volume was either 2 μL (induced leaf discs) or 10 μL (uninduced leaf discs, whole seedlings, and seedling roots). Data was visualized using DataAnalysis Version 4.3 (Bruker Compass DataAnalysis 4.3 (x64), Bruker Daltonik GmbH) and automated peak integration was performed with QuantAnalysis Version 4.3 (Bruker Compass DataAnalysis 4.3 (x64), Bruker Daltonik GmbH). Compounds were identified by comparison to commercial standards: anabasine [(±)-anabasine hydrochloride, Cayman Chemical, Ann Arbor, USA], anatabine [(*R*/*S*)-anatabine tartrate, Cayman Chemical, Ann Arbor, USA], caffeine (Sigma-Aldrich, St. Louis, USA), dihydrometanicotine (dihydrometanicotine dihydrochloride, Toronto Research Chemicals, Toronto Canada), and nicotine [(-)-nicotine, Fluka, Morristwon, USA]. Quantification of nicotine, anabasine and anatabine was carried out using external calibration curves.

### (*R*)- and (*S*)-nicotine analysis

Chiral LC-MS analysis was performed as indicated above (LC-MS analysis) with the following modifications. Separation was achieved on a Lux^®^ 3 μm AMP column (150 x 3.0 mm, Phenomenex, Torrance, USA) with an eluent flow rate of 0.3 mL/min and a column temperature of 40°C. Mobile phases A and B consisted of 10 mM ammonia bicarbonate (pH 9.2) in water and 2-propanol, respectively, and the elution program was isocratic at 20% B with a total run time of 20 min. Injection volumes were either 2 μL (MeJa-induced leaf discs) or 10 μL (uninduced leaf discs). Compounds were identified by comparison to commercial standards: caffeine (Sigma-Aldrich, St. Louis, USA), (-)-nicotine (Fluka, Morristwon, USA), and (±)-nicotine (Sigma-Aldrich, St. Louis, USA

## RESULTS

### An *N. benthamiana* genome assembled from Chromium linked reads and ONT-generated long reads

The use of Chromium linked-reads and ONT long reads resulted in a genome assembly of 2,995.0 Mb, consisting of 50,663 scaffolds, with an N50 of 14.0 Mb and a L50 of 67. The assembly has high completeness, as evidenced by a BUSCO completeness of 97.6% and a k-mer completeness of 97.82%. Additionally, the assembly quality value (QV) score of 37,88, indicates good per base accuracy. Comparison of our assembly with a recently published assembly generated using Pacific BioScience Highly Accurate Long Read Sequencing chemistry (PacBio HiFi) and HiC (Omni-C reads), indicates that our assembly is of comparable completeness and quality (Table S5). However, the HiFi+HiC assembly exhibits higher contiguity, as evidenced by the larger N50 and the smaller number of contigs. The assembly is available for download and can be accessed at the following link: https://opendata.earlham.ac.uk/opendata/data/Patron_2023-03-01_Nicotiana_benthamiana_genome_assembly/

### *N. benthamiana* encodes five *BBL* genes

Using the *NtBBL* genes as queries, we identified five candidate *BBL* genes in the *N. benthamiana* genome, four of which (*NbBBLa-d*) encoded open reading frames that translate to proteins of similar length to the BBL proteins encoded in the *N. tabacum* genome. The fifth gene, *NbBBLd’*, contained a premature stop codon (Figure 2A and Figure S2). Phylogenetic analysis indicates that *NbBBLa-d* may be orthologues of the equivalently named genes in *N. tabacum* but there is no obvious orthologue for *NtBBLe in the N. benthamiana* genome (Figure S2).

**Figure 2.**
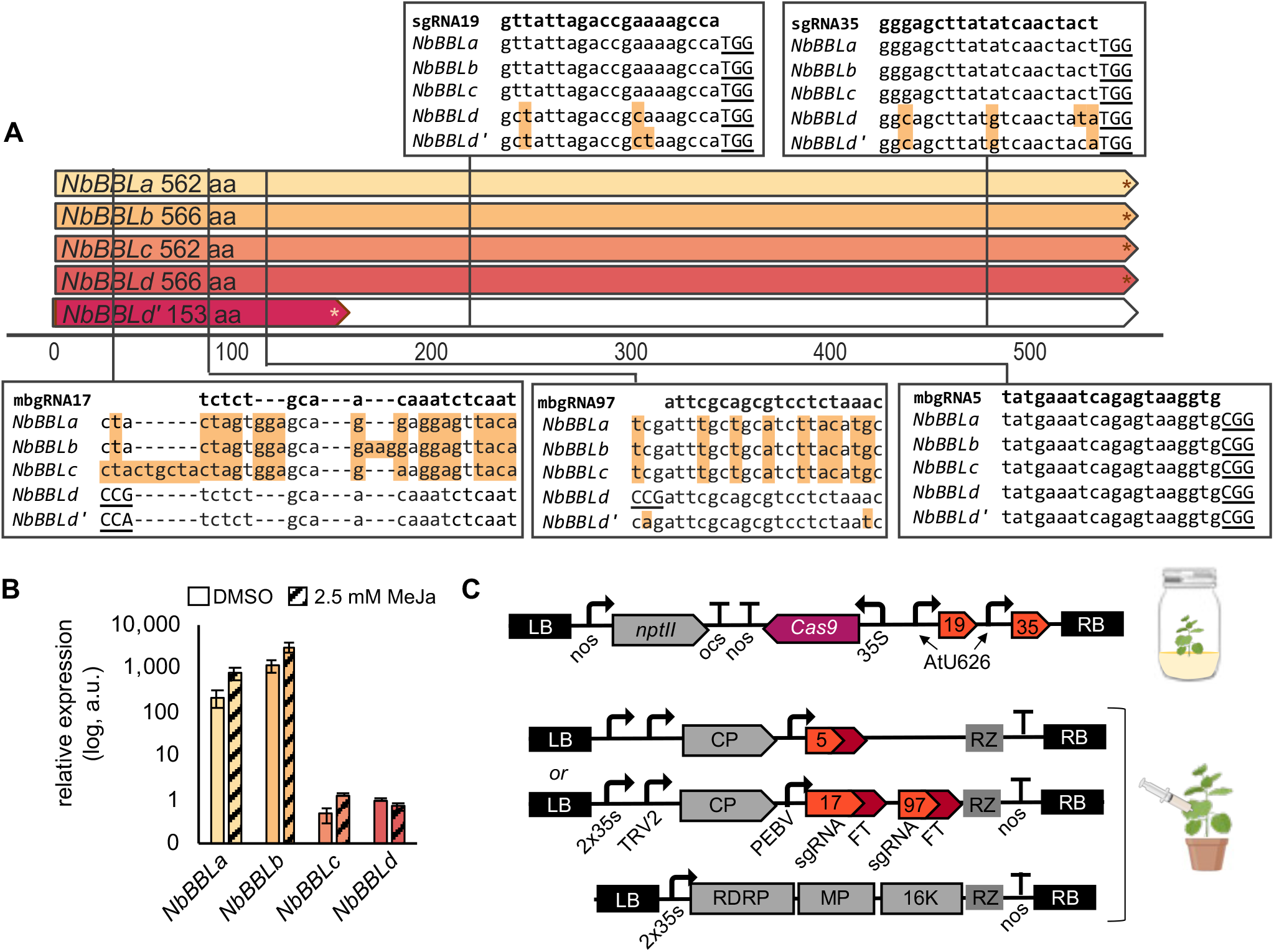
*Nicotiana benthamiana* berberine bridge enzyme-like (NbBBL) enzymes. **A.** Schematic of NbBBL enzymes indicating the locations and sequences of guide RNAs used for Cas9-mediated targeted mutagenesis. Canonical NGG protospacer adjacent motifs are underlined, * = stop codon. Background shading of bases indicates a mismatch to the guide RNA sequence. **B.** Expression of *NbBBL* genes relative to *NbEF1a* in roots of plants following treatment with 2.5 mM methyl jasmonate. Expression was determined by reverse transcription quantitative PCR (RT-qPCR). Error bars indicate the standard error of the mean of three biological replicates (4 technical replicates per sample). **C.** Schematics of plasmid constructs used for Cas9-mediated targeted mutagenesis by integration of T-DNA (above) or transient expression of a mobile guide RNA (mbgRNA). LB, left border; RB, right border; RZ, self-cleaving ribozyme; MP, movement protein, RDRP, RNA-dependent RNA polymerase; 16k, 16k gene; nos, nopaline synthase promoter or terminator; ocs, octopine synthase terminator; PEBV, pea early browning virus promoter; 35s, cauliflower mosaic virus 35s promoter; FT, flowering locus T mobile signal. This figure contains graphics from Biorender (biorender.com).

We analyzed the expression levels of *NbBBLa-d* in root tissues using reverse-transcription quantitative (realtime) polymerase chain reaction (RT-qPCR). *NbBBLa* and *NbBBLb* were expressed at a higher level than *NbBBLc* and *NbBBLd* (Figure 2B) and the expression of all genes except *NbBBLd* increased following exposure of plants to MeJa (Figure 2B).

### Production of *N. benthamiana* lines with Cas9-mediated targeted mutations in *NbBBL* genes

We first constructed a binary vector expressing two single guide RNAs (sgRNA19 and sgRNA35) for the stable transformation of wild-type *N. benthamiana* plants to introduce Cas9-mediated targeted mutations in *NbBBLa-c* (Figure 2A and C). The genotypes of transgenic (T_0_) plants were determined by PCR-amplification and sequencing of target loci. Lines in which the genotype was unclear or indicated genetic chimerism were discarded and T_1_ seed was collected from lines with homozygous, heterozygous or biallelic mutations in one or more *NbBBL* genes. The genotype at each locus as well as the presence or absence of the T-DNA was analyzed in T_1_ plants. T_2_ seeds were collected from plants in which the T-DNA had segregated and that contained homozygous or biallelic mutations at one or more *NbBBL* genes. Genotypes were confirmed in T2 plants and are provided in Figure 3 and Tables S6-S10. We also constructed two TMV-based viral vectors encoding mobile guide RNAs (mbgRNA) as previously described (Ellison *et al*., 2020). The first vector contained a single mbgRNA, mbgRNA5, targeting a single locus present in all *NbBBL* genes (Figure 2A and C). A second vector contained two mbgRNAs, mbgRNA17 and mbgRNA97, targeting sequences unique to *NbBBLd/d’* (Figure 2A and C). These vectors were infiltrated into the leaves of transgenic plants constitutively expressing Cas9, and progeny seeds were collected from pods that formed on stems in which mutations were detected at the target sites (see Methods). The genotypes of E_1_ plants were analyzed, and individuals with homozygous or biallelic mutations at loci of interest were selected (Figure 3, Tables S6-S10).

**Figure 3.**
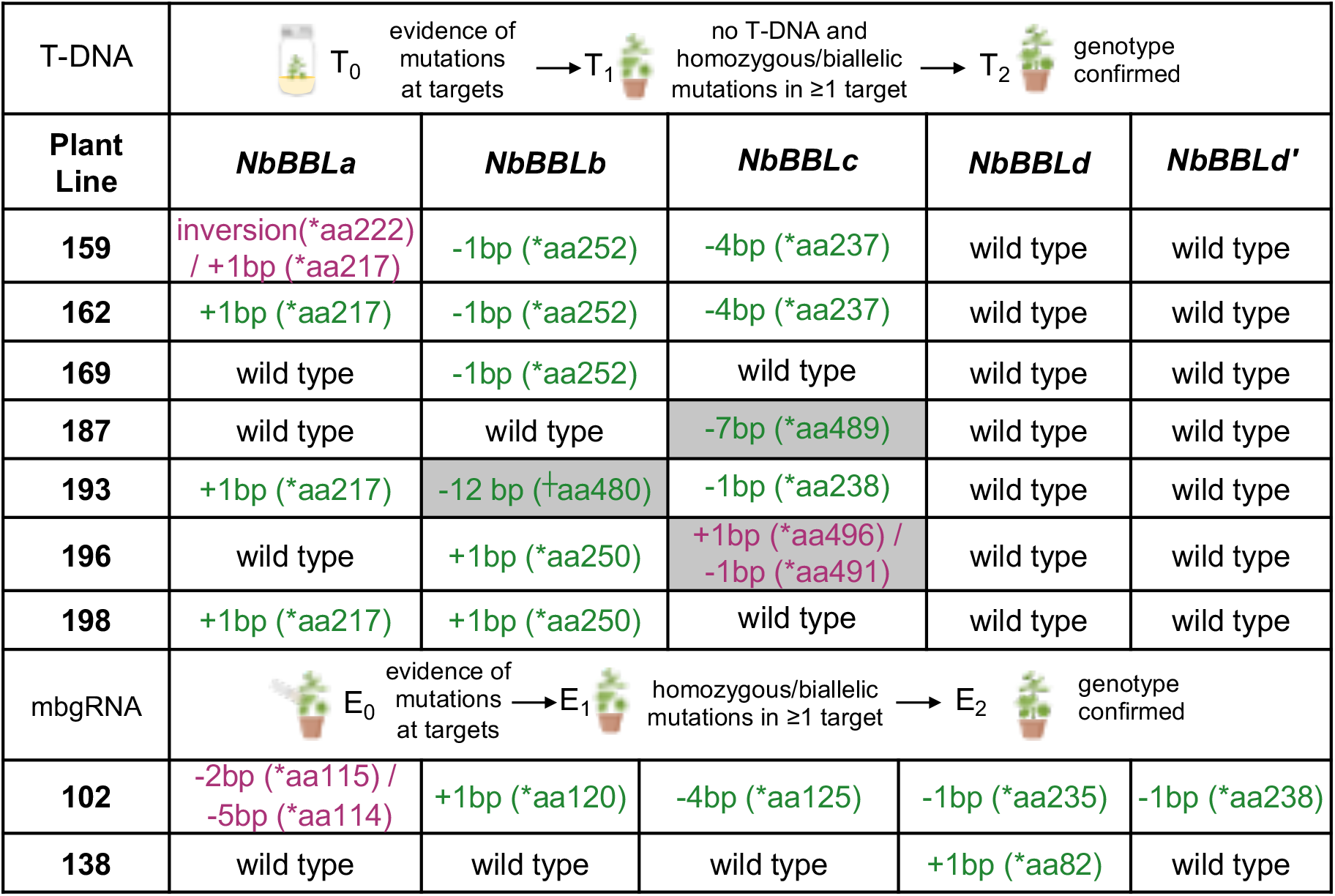
Workflow for the selection of gene-edited lines, and genotypes of all *NbBBL* genes in each line. The size of insertions (+) and deletions (-) are provided in base pairs (bp) together with position of the first premature stop codon (*aa). Where lines are mutated at two positions, only the first disruptive mutation is shown; details of all mutations can be found in Supplemental Data Tables S6-S10. Green font = homozygous mutation, purple font = biallelic mutation; gray shading = mutation may not result in loss of activity due to size, type or location, ┼ = in line 193, five residues of *NbBBLb* (AYINY) are replaced with a new residue (D). This figure contains graphics from Biorender (biorender.com).

Overall all guide RNAs except for mbgRNA17 resulted in mutations at the target locus. From these two methods, we were able to obtain lines with mutations in NbBBLb, NbBBLc or NbBBLd alone, as well as lines with mutations in combinations of two, three or five genes (Figure 3, Tables S6-S10).

### Inactivation of *NbBBL* genes leads to low pyridine alkaloid levels

We analyzed the levels of nicotine, anabasine, and anatabine in leaves of the different *NbBBL* mutant lines (T_2_/E_2_ generation). We included three different control lines: a wild type line (WT; control for all lines), a tissue culture control (TC WT; a wild type line recently regenerated in tissue culture, used as additional control for lines 159-198), and a transgenic Cas9 line (Cas9; the parental line constitutively expressing Cas9, used as additional control for lines 102 and 138) (Figure 4). In general, the levels of nicotine were always the highest, followed by anabasine and then anatabine. Single *NbBBLb*, *NbBBLc*, and *NbBBLd* mutants did not differ from control lines, and neither did the double *NbBBLb/c* mutant. Gratifyingly, the double *NbBBLa/b* mutant displayed lower nicotine levels compared to the control lines, although this difference was not significant in the absence of MeJa induction. Triple *NbBBLa/b/c* and quintuple *NbBBLa/b/c/d/d’* mutants did not appear to vary in nicotine content compared to the double *NbBBLa/b* mutant. However, one of the triple mutant lines (line 193) as well as the quintuple mutant displayed significantly lower nicotine content compared to the corresponding control lines under both induced and uninduced conditions (~3-4 fold reduction). The overall picture for anabasine and anatabine was similar. However, the anatabine levels in the triple and quintuple mutants were below our lowest calibration point and thus were not quantified (Figure 4).

**Figure 4.**
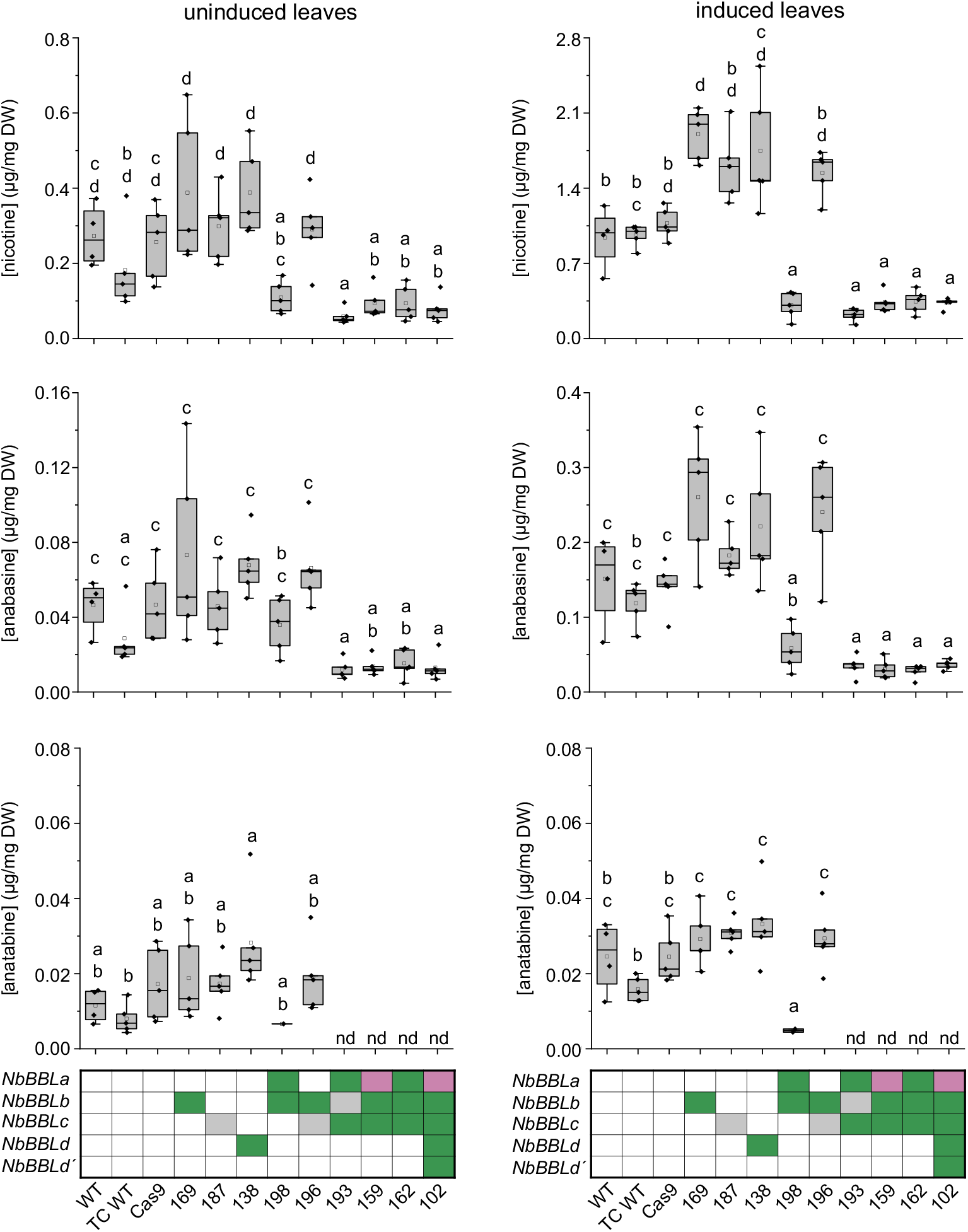
Pyridine alkaloid content in leaves of *N. benthamiana BBL* mutant in comparison to control lines as analyzed by LC-MS. Three control lines were analyzed: WT, wildtype (control for all lines); TC WT, tissue culture control (additional control for lines 159-198); Cas9, transgenic plants constitutively expressing Cas9 (additional control for lines 102 and 138). Graphs to the left represent the levels of nicotine, anabasine, and anatabine in leaves of greenhouse-grown plants (nd, not determined). Graphs to the right represent the analogous measurements 5 days after induction with 0.1% MeJa. For each line, four to five biological replicates were measured (black diamonds). Results are presented as box plots, with the center line representing the median, the unfilled square representing the mean, the box delimiting the upper and lower quartiles, and the whiskers representing the outlier range with a coefficient of 1.5. Significant differences were determined using ANOVA and *post hoc* Tukey tests on log transformed data. ANOVA p-values can be found in Table S11. The genotypes of *NbBBL* genes in each of the edited and control plants are indicated below the graphs, with green shading representing a homozygous knockout, purple shading indicating a biallelic knockout, and gray shading indicating uncertainty with respect to loss of function due to size, type or location of the mutation. Details of all mutations are provided in Figure 3 and Tables S6-S10.

### Inactivation of *NbBBL* genes leads to a racemic nicotine mixture

The analysis of pyridine alkaloids described above did not discriminate between *S* and *R* enantiomers. Thus, we switched to chiral LC-MS analysis to determine whether inactivation of *NbBBL* genes had any effect on enantiomeric purity.

We used a method able to separate (*S*)- and (*R*)- nicotine with near baseline resolution, and we focused on two control lines (wildtype and Cas9 transformant) versus the quintuple *NbBBLa/b/c/d/d’* mutant (line 102). While the control lines accumulated (*S*)-nicotine almost exclusively, the residual nicotine in the quintuple mutants was fully racemic (Figure 5, Figure S3).

**Figure 5.**
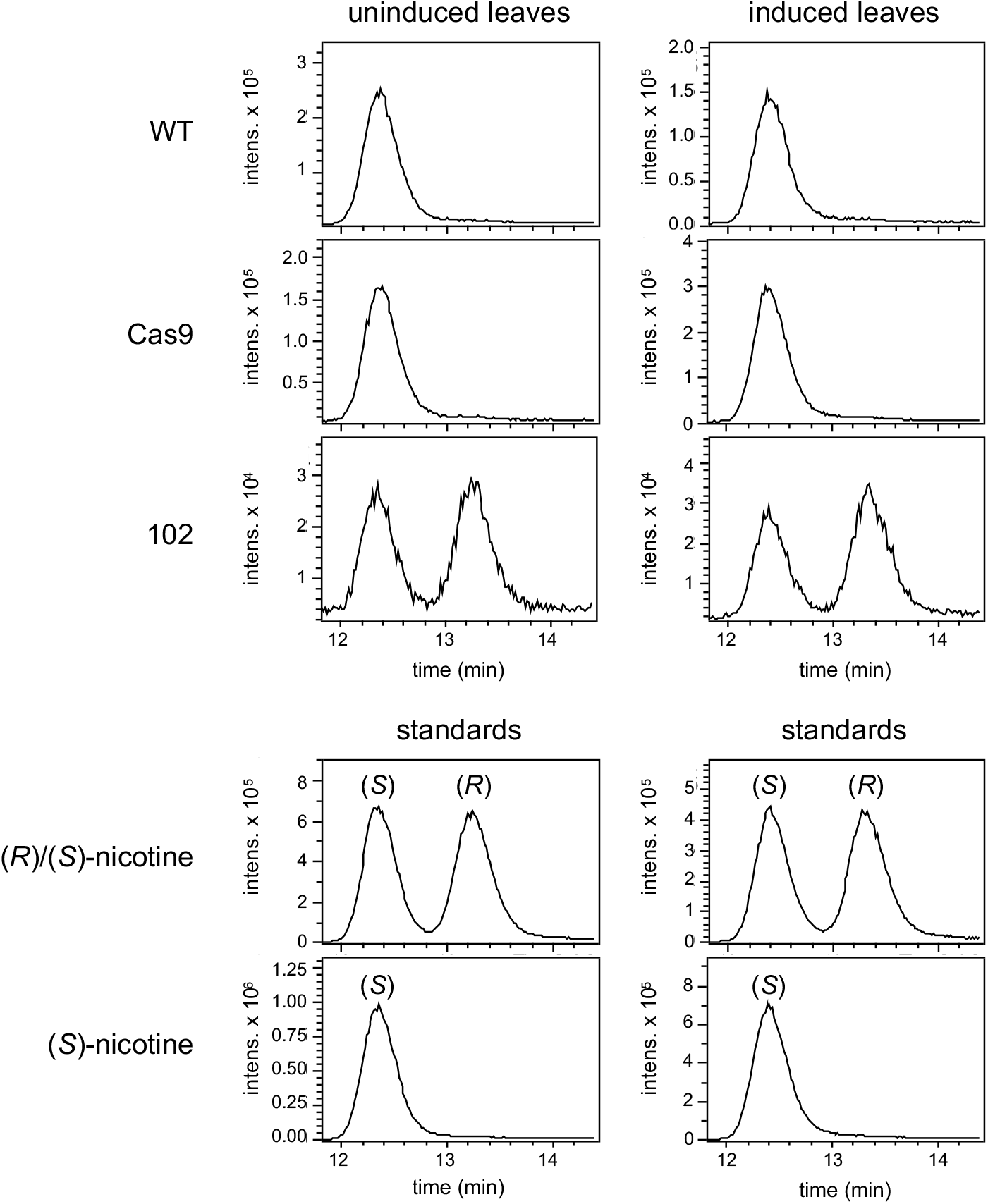
Analysis of (*S*)- and (*R*)-nicotine in leaves of the quintuple *NbBBL* mutant (line 102) in comparison to control lines (WT and Cas9). Traces correspond to extracted ion chromatograms (nicotine, [M+H]^+^) resulting from chiral LC-MS analyses (Lux^®^ 3 μm AMP column). Traces on the left are from uninduced plants, while traces on the right are from the same plants 5 days after induction with MeJa. Higher injection volumes were used for all uninduced samples (10 μL compared to 2 μL) to obtain comparable peak sizes. A total of four to five biological replicates were analyzed. While only one replicate is shown in this figure, all data is provided in Figure S3. The two bottom rows show the results of running a racemic nicotine standard and a (*S*)-nicotine standard along with the uninduced and induced samples.

### Inactivation of *NbBBL* genes prevents hydrogen loss from position C6

As mentioned in the introduction, feeding experiments with isotopically labeled nicotinic acid have shown the loss of hydrogen at position C6 during its incorporation into nicotine in *N. tabacum* (Dawson *et al*., 1960). Subsequent publications have supported this finding for nicotine biosynthesis (Leete & Liu, 1973) and extended it to anatabine and anabasine biosynthesis (Leete, 1978). Furthermore, nicotinamide has been found to incorporate into nicotine to a similar extent as nicotinic acid (Dawson *et al*., 1960).

We examined the incorporation of labeled D4-nicotinamide into nicotine, anabasine and anatabine in the two control lines (wildtype and Cas9 transformant) versus the quintuple *NbBBLa/b/c/d/d’* mutant, all lines under MeJa induction. We observed that the main labeled form of nicotine and anabasine in the fed control lines contained three deuterium atoms (D3 versions), demonstrating the loss of one deuterium atom during incorporation (Figure 6). By contrast, the quintuple mutant contained predominantly D4-nicotine and D4-anabasine instead of the corresponding D3 versions. For anatabine, in which both rings derive from the pyridine branch, we detected the D7 version predominantly, whereas D7 and D8 versions accumulated equally in the quintuple mutant (Figure 6). Interestingly, while only about 30% of nicotine was found labeled in these experiments, the percentage increased to approximately 50% in the case of anabasine, and around 80% in the case of anatabine (Figure 6).

**Figure 6.**
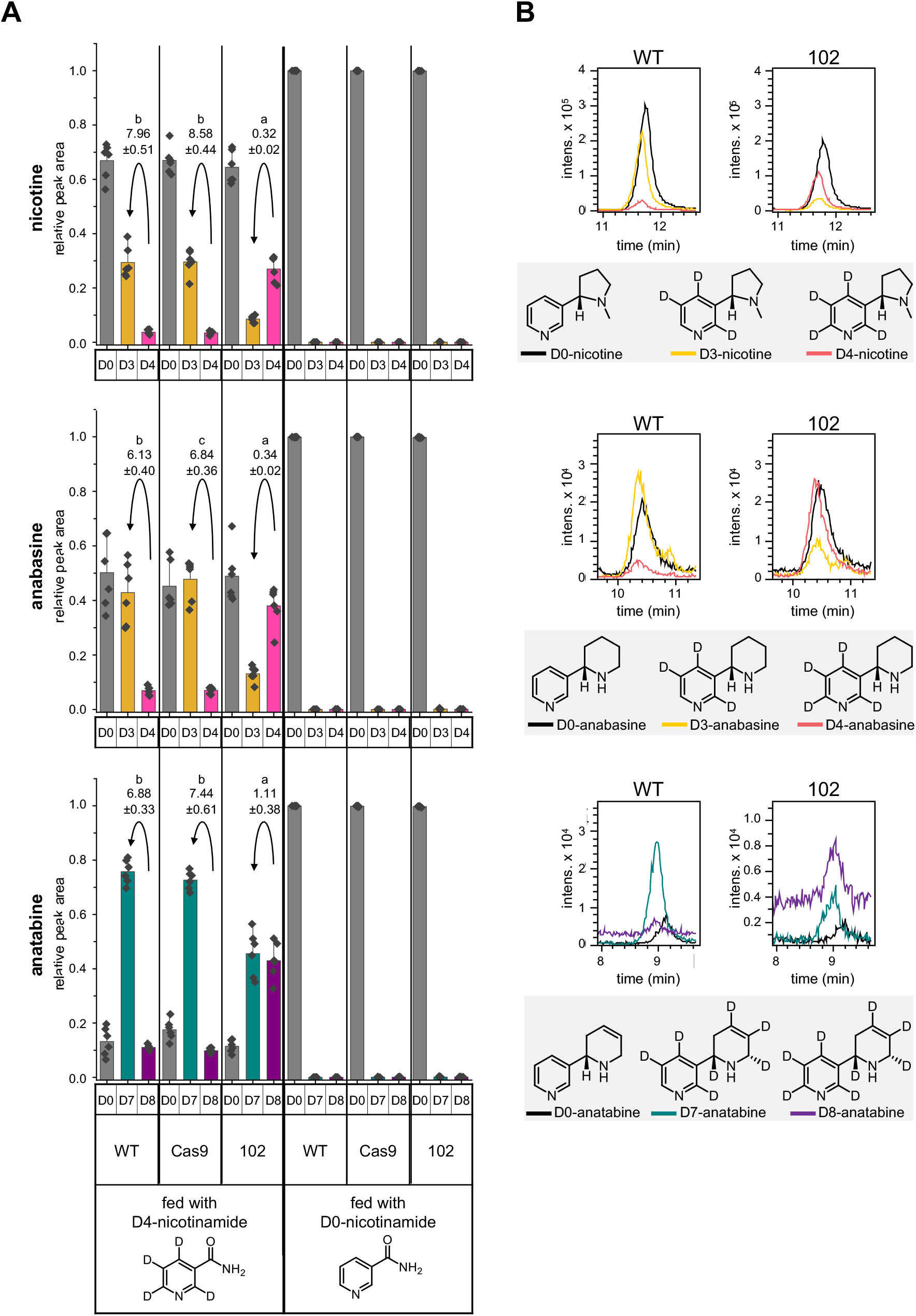
Incorporation of D4-nicotinamide into pyridine alkaloids in the quintuple *NbBBL* mutant (line 102) in comparison to control lines (WT and Cas9). Seedlings were grown under hydroponic conditions, induced with MeJa, and fed continuously for 5 days with either labeled (D4) or unlabeled (D0, control) nicotinamide via the roots. **A.** Relative levels of differently labeled nicotine, anabasine and anatabine in whole seedlings, as observed using LC-MS. For nicotine and anabasine, D0, D3, and D4 versions were observed and quantified. For anatabine, D0, D7, and D8 versions were observed and quantified. For each version, relative peak areas were obtained by dividing by the sum of peak areas of all labeled versions. Six biological replicates were analyzed (black diamonds). Bars represent mean values, and error bars represent the outlier range in plus direction with a coefficient of 1.5. Numbers above the arrows correspond to the fold-difference between D3 and D4 versions (for nicotine and anabasine) or between the D7 and D8 versions (for anatabine), plus/minus the standard deviation. Significant differences were determined using ANOVA and *post hoc* Tukey tests on log transformed (nicotine, anabasine) and untransformed (anatabine) data. ANOVA p-values can be found in Table S12. **B.** Representative traces (extracted ion chromatograms, [M+H]^+^) of the differently labeled pyridine alkaloids in quintuple *NbBBL* mutants (line 102) compared to WT plants. The colors of the depicted traces correspond to the colors of the corresponding bars in panel A. Chemical structures of each of the analyzed compounds are shown inside the gray boxes under each pair of graphs. The position of the lost deuterium atom was inferred from the comprehensive feeding studies carried out previously (Dawson *et al*., 1960; Leete & Liu, 1973; Leete, 1978).

### Inactivation of *NbBBL* genes leads to accumulation of DMN

Apart from displaying low nicotine levels, *N. tabacum* RNAi lines downregulated in *BBL* genes accumulate DMN in biosynthetic tissues (Kajikawa *et al*., 2011; Lewis *et al*., 2015). To examine whether a similar accumulation occurred in our gene-edited *N. benthamiana* plants, we harvested MeJa-induced roots from hydroponically grown quintuple mutants (line 102) and analyzed them via LC-MS. DMN was unequivocally detected in the mutant roots, while the compound was undetectable in the control lines (WT and Cas9) (Figure 7, Figure S4).

**Figure 7.**
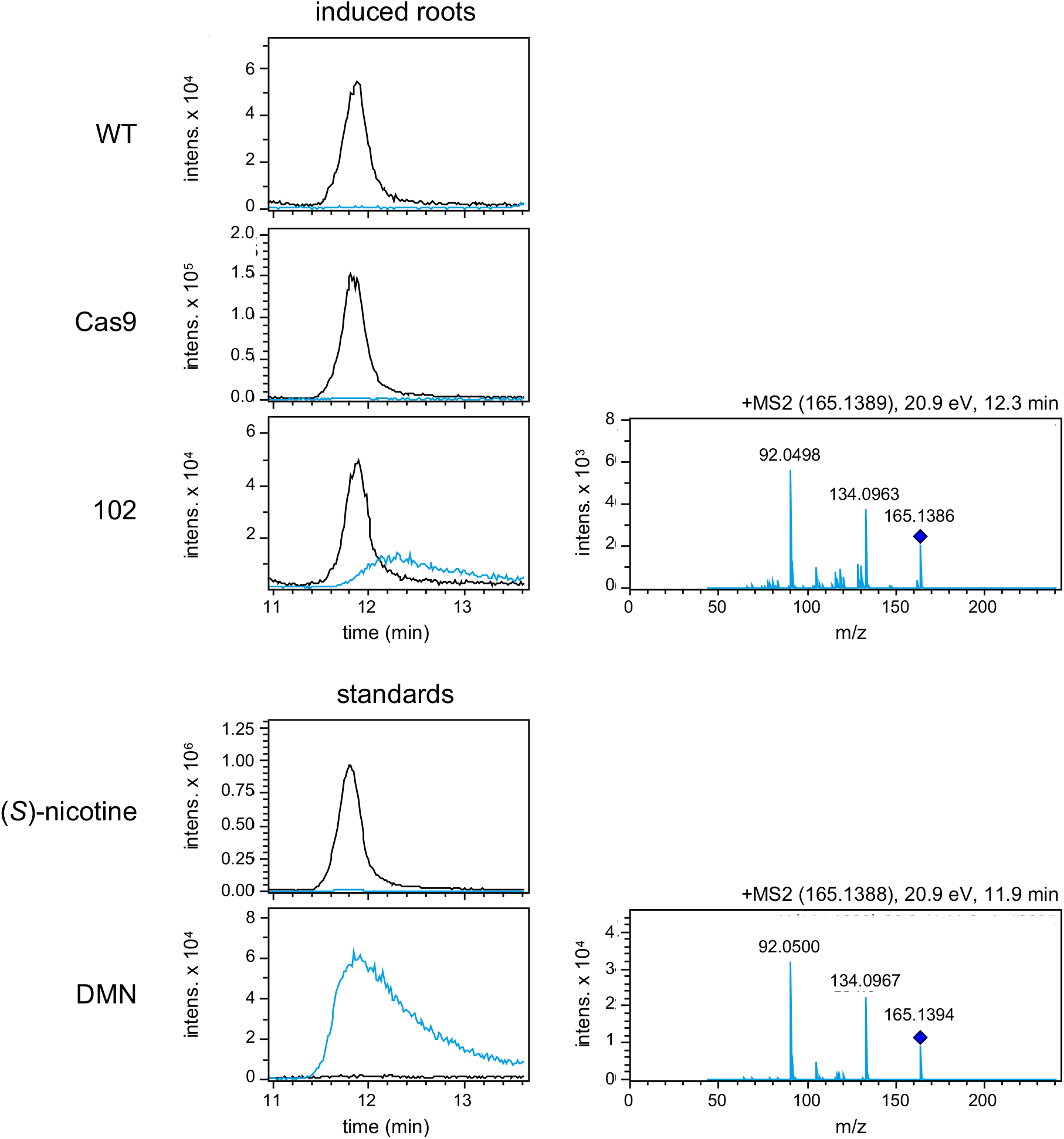
DMN accumulation in roots of the quintuple *NbBBL* mutant (line 102) as analyzed by LC-MS. Seedlings were grown under hydroponic conditions, and roots were harvested 5 days after induction with MeJa. Traces are extracted ion chromatograms ([M+H]^+^) corresponding to nicotine (black) or DMN (blue). Results from mutant line 102 are compared to results from two control lines (WT and Cas9). A total of five to six biological replicates per line were analyzed. While only one replicate is shown in this figure, all data is provided in Figure S4. Also shown are two traces corresponding to a (*S*)-nicotine standard and a DMN standard. In addition, MS2 spectra for DMN ([M+H]^+^) from line 102 and from the DMN standard are shown to the right. The labels at the top of the MS spectra indicate the mass of the fragmented ion (parenthesis), the collision energy, and the retention time.

## DISCUSSION

In this study we report the production and analysis of low-nicotine *N. benthamiana* lines using CRISPR/Cas9 to inactivate combinations of *BBL* genes, which are known to be involved in pyridine alkaloid biosynthesis. Our choice of *BBL* genes mirrors the choice taken by Lewis et al. when aiming at low-nicotine *N. tabacum* lines (Lewis *et al*., 2015). Briefly, aiming at genes in the pyrrolidine branch (e.g. *PMT*) is likely to result in lower nicotine levels at the expense of increased anatabine levels (Chintapakorn & Hamill, 2003; Wang *et al*., 2009). Conversely, aiming at genes involved only in the pyridine branch (e.g. *QPT*) may interfere with NAD metabolism (Lewis *et al*., 2015). *A622* and *BBL* genes would seem to be suitable targets based on their involvement in the late pathways towards the three pyridine alkaloids. However, Kajikawa et al. reported a failure to obtain *N. tabacum* lines constitutively silenced in *A622*, possibly due to the accumulation of nicotinic acid (Kajikawa *et al*., 2009). It should also be noted that two transcription factors represented by the *nic1* and *nic2* loci are known to control nicotine biosynthesis in *N. tabacum* (Shoji *et al*., 2010; Shoji & Hashimoto, 2012; Qin *et al*., 2021). These loci are not commonly used in tobacco breeding due to lower yields (Legg *et al*., 1970; Chaplin & Weeks, 1976) likely caused by overaccumulation of polyamines (Nölke *et al*., 2018). Thus, *BBL* genes seemed to be the best targets for our purposes in *N. benthamiana*.

Our first task was the identification of all *BBL* gene copies in *N. benthamiana*. At the start of our work, there were two published *N. benthamiana* genome drafts, both of which had identified several examples of homeologous genes consistent with an allotetraploid origin (Bombarely *et al*., 2012; Naim *et al*., 2012). However, these genome drafts were constructed from reads generated using Illumina-chemistry and, due to the complex nature of the genome, the assemblies consisted of more than 140,000 scaffolds. To obtain an improved draft, we generated a new *N. benthamiana* genome assembly using Chromium™ linked-read sequencing (10x Genomics) as well as Nanopore sequencing (Oxford Nanopore). The completeness of the assembly is comparable to a recently reported draft assembled from Pacific BioScience Highly Accurate Long Read Sequencing reads (PacBio HiFi) (Kurotani *et al*., 2023), however, the HiFi+HiC assembly has higher contiguity (Table S5). This comparison provides insights into the limitations and advantages of different sequencing technologies, which will inform future efforts in genome assembly and analysis and highlights the benefits of chromosome-spanning connectivity information. The availability of multiple long-read datasets provides the opportunity for further improvements by the community.

Out of the five *NbBBL* genes identified in the *N. benthamiana* genome, four were full-length (*NbBBLa*, *b*, *c*, and *d*), three were MeJa-responsive (*NbBBLa*, *b*, and *c*), and two were expressed at high levels (*NbBBLa* and *b*) (Figure 2A-B). We carried out gene editing of *NbBBL* genes using two different approaches. The first was a traditional approach targeting *NbBBLa, b*, and *c*, by integrating a T-DNA encoding a pair of sgRNAs to maximize the chances of inducing loss-of-function mutations (sgRNA19 and 35) (Figure 2A,C). We isolated a number of transgene-free lines with mutations at at least one of the two sgRNA sites, including a double *NbBBLa/b* knockout (line 198) and at least two triple *NbBBLa/b/c* knockouts (lines 159 and 162) (Figure 3). For the second gene-editing approach, we used virus-delivered mobile guide RNAs (mbgRNAs) (Ellison *et al*., 2020) which, in one embodiment, targeted all *BBL* genes (mbgRNA5) (Figure 2A,C). With this strategy, we succeeded in obtaining a quintuple knockout (line 102).

Metabolite analysis in leaves of the mutant lines revealed that simultaneous inactivation of *NbBBLa* and *NbBBLb* is sufficient to lower the levels of pyridine alkaloids, at least under induced conditions (Figure 4). Further inactivation of *NbBBLc, NbBBLd* and *NbBBLd’* did not result in significantly lower levels of pyridine alkaloids (Figure 4). Together with the abovementioned gene expression results (Figure 2B), this indicates that *NbBBLa* and *NbBBLb* are the main *NbBBL* genes involved in pyridine alkaloid biosynthesis in *N. benthamiana*. A major role of *BBLa* and *BBLb* was also observed in *N. tabacum* (Lewis *et al*., 2015, 2020; Kajikawa *et al*., 2017). Indeed, Kajikawa and coworkers (2017) reported that *NtBBLa* and *NtBBLb* are by far the highest expressed *NtBBLs* in cultivar TN90. Consistent with this, Lewis *et al*. (2015) observed that double *NtBBLa/b* and triple *NtBBLa/b/c* knockouts (EMS-induced) in cultivar DH98-325-6 displayed comparably reduced levels of nicotine, nornicotine, and anatabine. A follow-up publication by Lewis et al. (2020) supported these findings by transferring the mutant alleles to different backgrounds (cultivars K326, TN 90 and TN90 SRC). Similar to our results with the quintuple *N. benthamiana* knockout, Lewis et al. did not observe further pyridine alkaloid reductions by inactivating the remaining *NtBBL* genes (*d1*, *d2* and *e*) on top of the triple *NtBBLa/b/c* mutant (TN90 SRC background).

In the light of both our study and the one by Lewis et al. (2020), it is puzzling that the multiple *NtBBL* knockout reported in a brief communication by Schachtsiek & Stehle (2019) did not accumulate any detectable nicotine. We set out to investigate the origin of the residual nicotine in our *NbBBL* knockouts. Chiral analysis revealed that, while control lines accumulated (*S*)-nicotine almost exclusively, the residual nicotine in our quintuple *NbBBL* knockout was fully racemic (Figure 5). Not only does this suggest that BBLs are fully responsible for the chirality of nicotine, it also suggests that the residual nicotine derives from a non-stereoselective, spontaneous reaction. In addition, we carried out precursor feeding studies with deuterated (D4) nicotinamide. In control plants, we observed the loss of one deuterium atom, leading to preferential accumulation of D3 compared to D4 pyridine alkaloid versions (Figure 6). This preferential accumulation was lost in the quintuple *NbBBL* knockout (Figure 6), strongly supporting the hypothesis of a spontaneous reaction. It is well known that, in biological systems, spontaneous reactions are sometimes made faster and/or more specific via the action of enzymes. In particular, some of the scaffold-forming reactions in plant alkaloid biosynthesis are subject to enzymatic control despite their spontaneous nature, as discussed recently by Lichman (2021). One example is the coupling of tryptamine and secologanin in the biosynthesis of monoterpene indole alkaloids. In this case, the spontaneous reaction is slow and gives rise to two stereoisomers, whereas the enzymatic reaction is stereoselective, giving rise only to the *S* product (Stöckigt & Zenk, 1977). In a similar fashion, we postulate that BBLs not only accelerate an otherwise spontaneous reaction, they also ensure exclusive accumulation of the *S* stereoisomer of nicotine, which is the more biologically active one (Yildiz, 2004).

One outstanding question remains: what is the precise reaction catalyzed by BBL enzymes? As mentioned before, localized accumulation of DMN has been observed in *N. tabacum* roots upon down-regulation or inactivation of *NtBBL* genes (Kajikawa *et al*., 2011; Lewis *et al*., 2015). We observed a similar accumulation of DMN in our quintuple *NbBBL* mutant, confirming that the catalytic role of BBLs is conserved between *N. tabacum* and *N. benthamiana*. Given that DMN does not appear to be a substrate and that it likely derives from an intermediate downstream of the coupling step, Kajikawa et al. (2011) proposed that BBLs catalyze the oxidation of an already coupled intermediate. This proposal is fully consistent with our precursor feeding experiments, which showed a BBL-dependent loss of hydrogen from the pyridine ring. It is also consistent with several known BBL-catalyzed reactions involving hydride removal next to a nitrogen atom (Daniel *et al*., 2017). However, the proposal seems to be at odds with the observed BBL-dependent creation of the *S* stereocenter in the pyrrolidine ring. The inconsistency stems from the fact that the stereocenter is likely established during the coupling. To bridge this gap, we propose a double function where BBLs catalyze the stereoselective coupling as well as the subsequent oxidation, as shown in Figure 8A.

**Figure 8.**
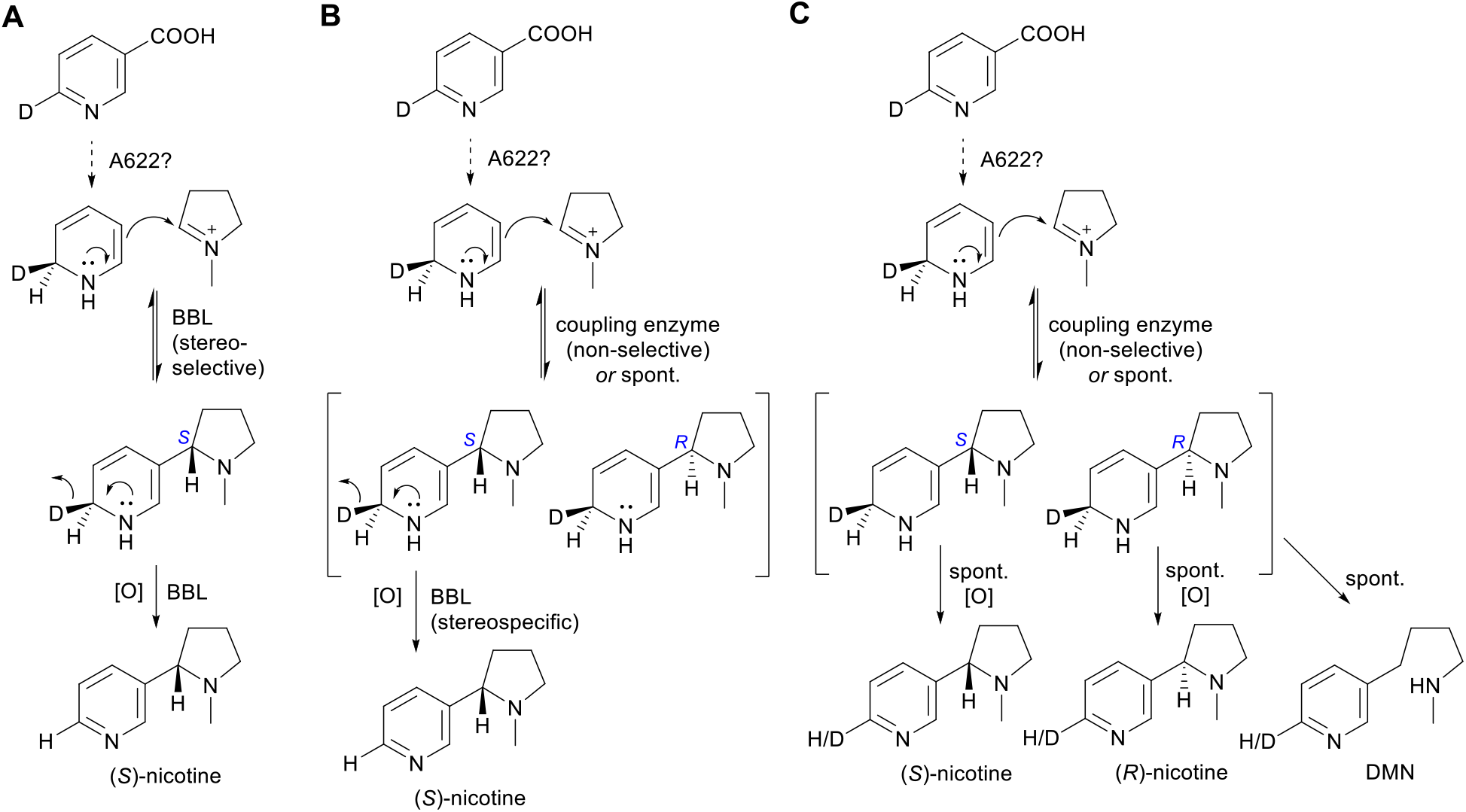
Mechanistic proposals for the BBL-catalyzed reaction(s). The proposals are based on our experimental results with the quintuple *NbBBL* knockout as well as previous results by Dawson and Leete (Dawson *et al*., 1960; Leete & Liu, 1973; Leete, 1978). A deuterium atom is shown at the C6 position in nicotinic acid (top molecule) to facilitate visualization of the fate of a hydrogen isotope at this position during biosynthesis. The stereochemistry of the initial reduction was established by Leete (Leete, 1978). In the BBL-catalyzed oxidations, the deuteride acceptor is likely to be FAD (not shown). [O], oxidation; spont., spontaneous; DMN, dihydrometanicotine. **A**. Scenario in which BBL enzymes catalyze both a stereoselective coupling and a subsequent oxidation. **B**. Alternative scenario in which the coupling is not stereoselective and is either spontaneous or catalyzed by an unknown enzyme. In this scenario, BBL enzymes are stereospecific for the *S* intermediate, thus giving rise only to (*S*)-nicotine. **C**. Proposal for the pyridine alkaloid pathway in the absence of functional BBL enzymes.

Yet, stereochemical control of the coupling reaction is not the only way BBL enzymes can be envisioned to direct nicotine biosynthesis towards the *S* form. In case of a non-BBL-catalyzed, non-stereoselective coupling leading to both *S* and *R* intermediates, BBL enzymes would still produce only (*S*)-nicotine if they were specific for the *S* intermediate. We show this alternative proposal in Figure 8B. In this scenario, an *R* intermediate would be produced, but it would be in fast equilibrium with the non-coupled forms, thus eventually converting to (*S*)-nicotine via decoupling.

Both of these proposals are consistent with our results, especially in light of the coupling reaction having an important degree of spontaneous character and the coupled intermediates being unstable. Upon inactivation of BBL enzymes as in our quintuple *NbBBL* mutant, the coupling would still occur (either spontaneously or catalyzed by an unknown enzyme), leading to production of both *S* and *R* intermediates. Apart from decoupling (reverse reaction), each coupled intermediate would have two possible fates: either oxidize spontaneously to the corresponding nicotine form, or undergo ring-opening to give DMN (Figure 8C). In all of these cases, the loss of a hydrogen from position C6 would be non-specific, precluding the preferential accumulation of D3 forms in the case of feeding with D4-nicotinamide. Moreover, if hydrogen loss during oxidation was rate limiting, D3 forms of the pyrimidine alkaloids would predominate due to the kinetic isotope effect, and this is indeed what we observed for nicotine and anabasine (Figure 7).

If the reaction(s) catalyzed by BBLs can indeed occur spontaneously, no further reduction in pyridine alkaloids can be expected by targeting this pathway step. A promising approach for further reduction could be the stacking of mutations in the Nic1/Nic2 loci as described in *N. tabacum* by Lewis 2020. Indeed, combining their *NtBBL*a/b/c alleles with the existing *nic2* locus resulted in a line with lower nicotine levels, and these were further reduced when *nic1* was stacked on top (*NtBBLa/b/c/nic1/nic2* line) (Lewis *et al*., 2020). As mentioned before the *nic1/nic2* loci are associated with reduced leaf quality in tobacco. This trait is associated with delayed senescence, which is problematic for the tobacco industry, but may be acceptable and even desirable in the case of *N. benthamiana*. However, the increased levels of polyamines observed in *N. tabacum* (Nölke et al., 2018) would be undesirable for high-value compound production in *N. benthamiana*.

In summary, we present an improved *N. benthamiana* genome draft as well as low-nicotine lines edited in *NbBBL* genes. Further analysis of edited lines provided insights into the nature of the reaction catalyzed by BBLs and opened the possibility that BBLs are the sought-after coupling enzymes in pyridine alkaloid biosynthesis.

## Supporting Information

Table S1. Primers used for expression analysis of NbBBL genes

Table S2. Primers used for amplification of sgRNA scaffolds.

Table S3. Primers used for construction of the mobile single guide RNA plasmid vectors.

Table S4. Primers used for genotyping plants with Cas9-induced mutations.

Table S5. Comparison of *Nicotiana benthamiana* genome assemblies.

Table S6. NbBBLa genotypes in plants with Cas9-mediated mutations.

Table S7. NbBBLb genotypes in plants with Cas9-mediated mutations.

Table S8. NbBBLc genotypes in plants with Cas9-mediated mutations.

Table S9. NbBBLd genotypes in plants with Cas9-mediated mutations.

Table S10. NbBBLd’ genotypes in plants with Cas9-mediated mutations.

Table S11. ANOVA p-values for Figure 4. Table S12. ANOVA p-values for Figure 6.

Figure S1. Schematic and photographs of the hydroponic system.

Figure S2. Maximum likelihood tree and multiple sequence alignment of *N. tabacum* and *N. benthamiana BBL* genes.

Figure S3. Analysis of (*S*)- and (*R*)-nicotine in leaves of the quintuple *NbBBL* mutant (line 102) in comparison to control lines (WT and Cas9).

Figure S4. DMN accumulation in roots of the quintuple *NbBBL* mutant (line 102) in comparison to two control lines (WT and Cas9), as analyzed by LC-MS.

## Acknowledgements

Plasmids pNJB069 (pTRV1), pEE393, and pEE515 along with seeds of *N. benthamiana* expressing SpCas9 were a generous gift from Dan Voytas. We thank Oleg Raitskin for advice on the design and construction of gene editing vectors. We thank Wilfried Haerty for his invaluable advice and management of the genome assembly. We also thank the John Innes Centre Horticultural services team for help with plant husbandry and acknowledge the contributions of the Genomics Pipelines and Core Bioinformatics Group at the Earlham Institute.

## Data Availability

The assembled genome is available at: https://opendata.earlham.ac.uk/opendata/data/Patron_2023-03-01_Nicotiana_benthamiana_genome_assembly. Reads are available under accession numbers: ERR3971933, ERR3971934, ERR3971506, ERR3971507, ERR3971508, ERR3971509, ERR3972081, ERR3972082, ERR3972083, ERR3972084, ERR3972085, and ERX10379414

## Author contributions

TY, QMD, FGF, and NJP conceptualized the study. QMD and DH extracted genomic DNA and performed nanopore sequencing. MOF, GS and DS assembled the genome. QMD and TO performed gene expression experiments. TY designed and assembled plasmid constructs for stable transformation. QMD designed, assembled and delivered viral vectors for mutagenesis. MC and MAS performed all plant tissue culture with supervision from WAH. QMD and KV performed genotyping of edited plants. KV and DM performed the analysis of alkaloid content. KV conducted the precursor feeding experiments and investigated the stereochemistry of nicotine and the accumulation of DMN. FGF and NP were responsible for supervision and funding acquisition. KV, QMD, MOF, FGF and NP drafted the text and figures. All authors contributed to revising and editing the text.

## References

Armstrong DW, Wang X, Ercal N. 1998. Enantiomeric composition of nicotine in smokeless tobacco, medicinal products, and commercial reagents. Chirality 10: 587–591.

Armstrong DW, Wang X, Lee J-T, Liu Y-S. 1999. Enantiomeric composition of nornicotine, anatabine, and anabasine in tobacco. Chirality 11: 82–84.

Bally J, Jung H, Mortimer C, Naim F, Philips JG, Hellens R, Bombarely A, Goodin MM, Waterhouse PM. 2018. The Rise and Rise of *Nicotiana* benthamiana: A Plant for All Reasons. Annual Review of Phytopathology 56: 405–426.

Bally J, Nakasugi K, Jia F, Jung H, Ho SYW, Wong M, Paul CM, Naim F, Wood CC, Crowhurst RN, et al. 2015. The extremophile *Nicotiana benthamiana* has traded viral defence for early vigour. Nature Plants. 1: 15165.

Bombarely A, Rosli HG, Vrebalov J, Moffett P, Mueller LA, Martin GB. 2012. A Draft Genome Sequence of Nicotiana benthamiana to Enhance Molecular Plant-Microbe Biology Research. Molecular Plant-microbe Interactions: MPMI 25: 1523–1530.

Brückner K, Tissier A. 2013. High-level diterpene production by transient expression in *Nicotiana benthamiana*. Plant Methods 9: 46.

Chaplin JF, Weeks WW. 1976. Association between percent total alkaloids and other traits in flue-cured tobacco 1. Crop Science 16: 416–418.

Chen B, Gilbert L a., Cimini B a., Schnitzbauer J, Zhang W, Li G-W, Park J, Blackburn EH, Weissman JS, Qi LS, et al. 2013a. Dynamic imaging of genomic loci in living human cells by an optimized CRISPR/Cas system. Cell 155: 1479–1491.

Chen Q, Lai H, Hurtado J, Stahnke J, Leuzinger K, Dent M. 2013b. Agroinfiltration as an Effective and Scalable Strategy of Gene Delivery for Production of Pharmaceutical Proteins. Advanced Techniques in Biology & Medicine 1: 103.

Chintapakorn Y, Hamill JD. 2003. Antisense-mediated down-regulation of putrescine N-methyltransferase activity in transgenic *Nicotiana tabacum* L. can lead to elevated levels of anatabine at the expense of nicotine. Plant Molecular Biology 53: 87–105.

Daniel B, Konrad B, Toplak M, Lahham M, Messenlehner J, Winkler A, Macheroux P. 2017. The family of berberine bridge enzyme-like enzymes: A treasure-trove of oxidative reactions. Archives of Biochemistry and Biophysics 632: 88–103.

D’Aoust MA, Couture MMJ, Charland N, Trépanier S, Landry N, Ors F, Vézina LP. 2010. The production of hemagglutinin-based virus-like particles in plants: A rapid, efficient and safe response to pandemic influenza. Plant Biotechnology Journal 8: 607–619.

Davis K, Gkotsi DS, Smith DRM, Goss RJM, Caputi L, O’Connor SE. 2020. Nicotiana benthamiana as a Transient Expression Host to Produce Auxin Analogs. Frontiers in plant science 11: 581675.

Dawson RF, Christman DR, D’Adamo A, Solt ML, Wolf AP. 1960. The Biosynthesis of Nicotine from Isotopically Labeled Nicotinic Acids1. Journal of the American Chemical Society 82: 2628–2633.

Dong L, Jongedijk E, Bouwmeester H, Van Der Krol A. 2016. Monoterpene biosynthesis potential of plant subcellular compartments. The New Phytologist 209: 679–690.

Dudley QM, Jo S, Guerrero DAS, Chhetry M, Smedley MA, Harwood WA, Sherden NH, O’Connor SE, Caputi L, Patron NJ. 2022a. Reconstitution of monoterpene indole alkaloid biosynthesis in genome engineered *Nicotiana benthamiana*. Communications biology 5: 949.

Dudley QM, Raitskin O, Patron NJ. 2022b. Cas9-Mediated Targeted Mutagenesis in Plants. Methods in Molecular Biology 2379: 1–26.

Ellison EE, Nagalakshmi U, Gamo ME, Huang P-J, Dinesh-Kumar S, Voytas DF. 2020. Multiplexed heritable gene editing using RNA viruses and mobile single guide RNAs. Nature Plants 6: 620–624.

Engler C, Youles M, Grüetzner R. 2014. A Golden Gate modular cloning toolbox for plants. ACS Synthetic Biology 3: 839–843.

Forman V, Luo D, Geu-Flores F, Lemcke R, Nelson DR, Kampranis SC, Staerk D, Møller BL, Pateraki I. 2022. A gene cluster in *Ginkgo biloba* encodes unique multifunctional cytochrome P450s that initiate ginkgolide biosynthesis. Nature Communications 13: 5143.

Geu-Flores F, Nielsen MT, Nafisi M, Møldrup ME, Olsen CE, Motawia MS, Halkier BA. 2009. Glucosinolate engineering identifies a gamma-glutamyl peptidase. Nature Chemical Biology 5: 575–577.

Gnanasekaran T, Vavitsas K, Andersen-Ranberg J, Nielsen AZ, Olsen CE, Hamberger B, Jensen PE. 2015. Heterologous expression of the isopimaric acid pathway in *Nicotiana benthamiana* and the effect of N-terminal modifications of the involved cytochrome P450 enzyme. Journal of Biological Engineering 9: 24.

Goodin MM, Zaitlin D, Naidu RA, Lommel SA. 2008. Nicotiana benthamiana: Its History and Future as a Model for Plant–Pathogen Interactions. Molecular plantmicrobe interactions: MPMI 21: 1015–1026.

GRAS Notice 910 Thaumatin II produced in Nicotiana benthamiana. 2020. Food and Drug Administration.

Grosse-Holz F, Kelly S, Blaskowski S, Kaschani F, Kaiser M, van der Hoorn RAL. 2018. The transcriptome, extracellular proteome and active secretome of agroinfiltrated *Nicotiana benthamiana* uncover a large, diverse protease repertoire. Plant Biotechnology Journal 16: 1068–1084.

van Herpen TWJM, Cankar K, Nogueira M, Bosch D, Bouwmeester HJ, Beekwilder J. 2010. Nicotiana benthamiana as a production platform for artemisinin precursors. PloS one 5.

Holley G, Beyter D, Ingimundardottir H, Møller PL, Kristmundsdottir S, Eggertsson HP, Halldorsson BV. 2021. Ratatosk: hybrid error correction of long reads enables accurate variant calling and assembly. Genome Biology 22: 28.

Kajikawa M, Hirai N, Hashimoto T. 2009. A PIP-family protein is required for biosynthesis of tobacco alkaloids. Plant Molecular Biology 69: 287–298.

Kajikawa M, Shoji T, Kato A, Hashimoto T. 2011. Vacuole-localized berberine bridge enzyme-like proteins are required for a late step of nicotine biosynthesis in tobacco. Plant Physiology 155: 2010–2022.

Kajikawa M, Sierro N, Kawaguchi H, Bakaher N, Ivanov NV, Hashimoto T, Shoji T. 2017. Genomic Insights into the Evolution of the Nicotine Biosynthesis Pathway in Tobacco. Plant Physiology 174: 999–1011.

Karasov TL, Chae E, Herman JJ, Bergelson J. 2017. Mechanisms to Mitigate the Trade-Off between Growth and Defense. The Plant Cell 29: 666–680.

Kurotani K-I, Hirakawa H, Shirasawa K, Tanizawa Y, Nakamura Y, Isobe S, Notaguchi M. 2023. Genome Sequence and Analysis of *Nicotiana benthamiana*, the Model Plant for Interactions between Organisms. Plant & Cell Physiology 64: 248–257.

Lau W, Sattely ES. 2015. Six enzymes from mayapple that complete the biosynthetic pathway to the etoposide aglycone. Science 349: 1224–1228.

Leete E. 1978. Stereochemistry of the reduction of nicotinic acid when it serves as a precursor of anatabine. Journal of the Chemical Society. Chemical communications: 610–611.

Leete E, Liu Y-Y. 1973. Metabolism of [2-3H]-and [6-3H]-nicotinic acid in intact *Nicotiana tabacum* plants. Phytochemistry 12: 593–596.

Legg PD, Collins GB, Litton CC. 1970. Heterosis and combining ability in diallel crosses of Burley tobacco, *Nicotiana tabacum L*. 1. Crop Science 10: 705–707.

Lewis RS, Drake-Stowe KE, Heim C, Steede T, Smith W, Dewey RE. 2020. Genetic and Agronomic Analysis of Tobacco Genotypes Exhibiting Reduced Nicotine Accumulation Due to Induced Mutations in Berberine Bridge Like (BBL) Genes. Frontiers in Plant Science 11: 368.

Lewis RS, Lopez HO, Bowen SW, Andres KR, Steede WT, Dewey RE. 2015. Transgenic and mutation-based suppression of a berberine bridge enzyme-like (BBL) gene family reduces alkaloid content in field-grown tobacco. PloS one 10: e0117273.

Li H. 2018. Minimap2: pairwise alignment for nucleotide sequences. Bioinformatics 34: 3094–3100.

Lichman BR. 2021. The scaffold-forming steps of plant alkaloid biosynthesis. Natural Product Reports 38: 103–129.

Liu Q, Majdi M, Cankar K, Goedbloed M, Charnikhova T, Verstappen FWA, de Vos RCH, Beekwilder J, van der Krol S, Bouwmeester HJ. 2011. Reconstitution of the costunolide biosynthetic pathway in yeast and Nicotiana benthamiana. PloS one. 6: e23255

Liu D, Shi L, Han C, Yu J, Li D, Zhang Y. 2012. Validation of reference genes for gene expression studies in virus-infected Nicotiana benthamiana using quantitative real-time PCR. PloS one 7: e46451.

Lomonossoff GP, D’Aoust MA. 2016. Plant-produced biopharmaceuticals: A case of technical developments driving clinical deployment. Science 353: 1237–1240.

Michael TP, Jupe F, Bemm F, Motley ST, Sandoval JP, Lanz C, Loudet O, Weigel D, Ecker JR. 2018. High contiguity Arabidopsis thaliana genome assembly with a single nanopore flow cell. Nature Communications 9: 1–8.

Miettinen K, Dong L, Navrot N, Schneider T, Burlat V, Pollier J, Woittiez L, van der Krol S, Lugan R, Ilc T, et al. 2014. The seco-iridoid pathway from Catharanthus roseus. Nature Communications 5: 3606.

Naim F, Nakasugi K, Crowhurst RN, Hilario E, Zwart AB, Hellens RP, Taylor JM, Waterhouse PM, Wood CC. 2012. Advanced Engineering of Lipid Metabolism in Nicotiana benthamiana Using a Draft Genome and the V2 Viral Silencing-Suppressor Protein. PloS one._7: e52717.

Nett RS, Sattely ES. 2021. Total Biosynthesis of the Tubulin-Binding Alkaloid Colchicine. Journal of the American Chemical Society 143: 19454–19465.

Nölke G, Volke D, Chudobová I, Houdelet M, Lusso M, Frederick J, Adams A, Kudithipudi C, Warek U, Strickland JA, et al. 2018. Polyamines delay leaf maturation in low-alkaloid tobacco varieties. Plant direct 2: e00077.

Qin Q, Humphry M, Gilles T, Fisher A, Patra B, Singh SK, Li D, Yang S. 2021. *NIC1* cloning and gene editing generates low-nicotine tobacco plants. Plant Biotechnology Journal 19: 2150–2152.

Reed J, Stephenson MJ, Miettinen K, Brouwer B, Leveau A, Brett P, Goss RJM, Goossens A, O’Connell MA, Osbourn A. 2017. A translational synthetic biology platform for rapid access to gram-scale quantities of novel drug-like molecules. Metabolic Engineering 42: 185–193.

Rhie A, Walenz BP, Koren S, Phillippy AM. 2020. Merqury: reference-free quality, completeness, and phasing assessment for genome assemblies. Genome Biology 21: 245.

Schachtsiek J, Stehle F. 2019. Dataset on nicotine-free, nontransgenic tobacco (Nicotiana tabacum l.) edited by CRISPR-Cas9. Data in Brief 26: 104395.

Shoji T, Hashimoto T. 2011. Nicotine Biosynthesis. In: Plant Metabolism and Biotechnology. Chichester, UK: John Wiley & Sons, Ltd, 191–216.

Shoji T, Hashimoto T. 2012. DNA-binding and transcriptional activation properties of tobacco *NIC2-* locus ERF189 and related transcription factors. Plant Biotechnology 29: 35–42.

Shoji T, Kajikawa M, Hashimoto T. 2010. Clustered transcription factor genes regulate nicotine biosynthesis in tobacco. The Plant Cell 22: 3390–3409.

Simão FA, Waterhouse RM, Ioannidis P, Kriventseva EV, Zdobnov EM. 2015. BUSCO: Assessing genome assembly and annotation completeness with single-copy orthologs. Bioinformatics.

Sisson, S. 1990. Alkaloid composition of the Nicotiana species. Beitr. Tabakforsch. Int. 14: 327–339.

Stephenson MJ, Reed J, Patron NJ, Lomonossoff GP, Osbourn A. 2020. 6.11 - Engineering Tobacco for Plant Natural Product Production. In: Liu H-W (ben), Begley TP, eds. Comprehensive Natural Products III. Oxford: Elsevier, 244–262.

Steppuhn A, Gase K, Krock B, Halitschke R, Baldwin IT. 2004. Nicotine’s defensive function in nature. PloS Biology 2: E217.

Stöckigt J, Zenk MH. 1977. Isovincoside (strictosidine), the key intermediate in the enzymatic formation of indole alkaloids. FEBS letters 79: 233–237.

Wang P, Zeng J, Liang Z, Miao Z, Sun X, Tang K. 2009. Silencing of PMT expression caused a surge of anatabine accumulation in tobacco. Molecular Biology Reports 36: 2285–2289.

Weisenfeld NI, Kumar V, Shah P, Church DM, Jaffe DB. 2017. Direct determination of diploid genome sequences. Genome Research 27: 757–767.

Xu S, Brockmöller T, Navarro-Quezada A, Kuhl H, Gase K, Ling Z, Zhou W, Kreitzer C, Stanke M, Tang H, et al. 2017. Wild tobacco genomes reveal the evolution of nicotine biosynthesis. Proceedings of the National Academy of Sciences of the United States of America 114: 6133–6138.

Yildiz D. 2004. Nicotine, its metabolism and an overview of its biological effects. Toxicon: official journal of the International Society on Toxinology 43: 619–632.

Zenkner FF, Margis-Pinheiro M, Cagliari A. 2019. Nicotine Biosynthesis in Nicotiana: A Metabolic Overview. Tobacco Science 56: 1–9.

