## Supporting Information for "An improved *Nicotiana benthamiana* bioproduction chassis provides novel insights into nicotine biosynthesis"

Table S1. Primers used for expression analysis of NbBBL genes.
Table S2. Primers used for amplification of sgRNA scaffolds.
Table S3. Primers used for construction of the mobile single guide RNA plasmid vectors.
Table S4. Primers used for genotyping plants with Cas9-induced mutations.
Table S5. Comparison of *Nicotiana benthamiana* genome assemblies.
Table S6. NbBBLa genotypes in plants with Cas9-mediated mutations.
Table S7. NbBBLb genotypes in plants with Cas9-mediated mutations.
Table S8. NbBBLc genotypes in plants with Cas9-mediated mutations.
Table S9. NbBBLd genotypes in plants with Cas9-mediated mutations.
Table S10. NbBBLd' genotypes in plants with Cas9-mediated mutations.
Table S11. ANOVA p-values for Figure 4.
Table S12. ANOVA p-values for Figure 6.

Figure S1. Schematic and photographs of the hydroponic system.
Figure S2. Maximum likelihood tree and multiple sequence alignment of *N. tabacum* and *N. benthamiana BBL* genes.
Figure S3. Analysis of (*S*)- and (*R*)-nicotine in leaves of the quintuple *NbBBL* mutant (line 102) in comparison to control lines (WT and Cas9).
Figure S4. DMN accumulation in roots of the quintuple *NbBBL* mutant (line 102) in comparison to two control lines (WT and Cas9), as analyzed by LC-MS.

**Table S1. Primers used for expression analysis of *NbBBL* genes.**

| | RTqPCR primer (forward) | RTqPCR primer (reverse) |
|---|---|---|
| **NbBBLa** | CGTGGTTCAACGCAAGAGAAC | ACTAACAACGGAATCTCTCTCAAGG |
| **NbBBLb** | TGATCCACTAAATGTTTTCCGC | ACAACGGAATCTCTCTCAACCT |
| **NbBBLc** | GTTCAACGCAAGACAATAAGTATAGC | CTCGTTGAATTGGATCAATATCGC |
| **NbBBLd** | CCTCAATGCAAGAGCATACCTAC | CCTTTATGATCCCAGGAGGCA |
| **NbEF1a** | AGCTTTACCTCCCAAGTCATC | AGAACGCCTGTCAATCTTGG |

**Table S2. Primers used for amplification of sgRNA scaffolds.** The plasmid vector pEPOR1CB0022 (Addgene#117537), which contains the sgRNA stem extension scaffold sequence first reported by Chen *et al. Cell* 155(7):1479–91 (2013), was used as a template. Details of the U6-promoter and L1 acceptor that the resulting amplicon was assembled with are also provided.

| sgRNA # | L1 acceptor | U6 Promoter | F primer | R primer |
|---|---|---|---|---|
| **19** | Position 3 pICH47751 Addgene #48002 | pICSL90002 (AtU6-26) Addgene #68261 | tgtGGTCTCtattGTTATTAGACCGAAAAGCCAgtttaagagctatgctggaaac | tGGTCTCtagcgaaaaaaagcaccgact |
| **35** | Position 4 pICH47761 Addgene #48003 | pICSL90002 (AtU6-26) Addgene #68261 | tgtGGTCTCtattGGGAGCTTATATCAACTACTgtttaagagctatgctggaaac | |

**Table S3. Primers used for construction of mobile single guide RNA plasmid vectors.** The plasmid vector pEPQDKN0761 (Addgene #185630) containing an sgRNA fused to truncated flowering locus T was used as a PCR template.

| | F Primer | R Primer |
|---|---|---|
| Primers to create an amplicon encoding mbgRNA5 used to construct a vector containing one mobile guide RNA targeting BBLa/BBLb/BBLc/BBLd/BBLd' (used to create plant line 102) | `aCGTCTCgcaggcacctgcaacgAAACTATGAAATCAGAGTAAGGTGgttttagagctag` | `tCGTCTCccgagcacctgctagtCACTttggccataagtaaccttt` |
| Primers to create an amplicon encoding mbgRNA17 used to construct a vector containing two mobile guide RNAs targeting BBLd and BBLd' (used to create plant line 138) | `aCGTCTCgcaggcacctgcaacgAAACATTGAGATTTGTTGCAGAGAgttttagagctag` | `tCGTCTCccgagtcacctgctagtGCACgctcaacacgtacccggccgcgattggccataagtaacctttagagt` |
| Primers to create an amplicon encoding mbgRNA97 used to construct a vector containing two mobile guide RNAs targeting BBLd and BBLd' (used to create plant line 138) | `aCGTCTCgcaggcacctgcaacgGTGCGTTTAGAGGACGCTGCGAATgttttagagctag` | `tCGTCTCccgagcacctgctagtCACTttggccataagtaaccttt` |

**Table S4. Primers used for genotyping plants with Cas9-induced mutations.** Asterisks indicate a phosphorothioate bond.

| | **PCR/sequencing primer (forward)** | **PCR/sequencing primer (reverse)** |
|---|---|---|
| **NbBBLa** | ttagatctcaaactgtatattatatatacaatgcag | ctaacaacggaatctctctcaag*g |
| **NbBBLb** | tattatccttctgttctcaaactgctc | ctaacaacggaatctctctcaac*c |
| **NbBBLc** | atgtttctactcataattctgatcagct | tttattaggaaagttaacaacgtacatttg |
| **NbBBLd** | ctgtgcactcttccttgtataaagc | gctgaaaatccatgaacgtc*g |
| **NbBBLd'** | gaagtaaacgaaagttgcaatgaa*g | aattaaagtaataggacaactaacaatatgacc |
| **Sequencing primer** Used to sequence amplicons of BBLa/BBLb/BBLc created using the primers above | aatttattacgccattgccaag | |
| | **ddPCR primer (forward)** | **ddPCR primer (reverse)** |
| ***nptII*** | cctgccgagaaagtatccat | tcttcgtccagatcatcctg |
| ***Rdr1*** **(reference gene)** | gttacgccatccatgtgttg | cagagttcaatttgccagca |

**Table S5. Comparison of *Nicotiana benthamiana* genome assemblies.** Metrics for the assemblies produced in this study (10x, 10x + ONT) and a recently published assembly (PacBio HiFI + HiC; Kurotani *et al.*, 2023). For the merqury k-mer analysis, two k-mer databases were constructed: one from the Illumina pair-end data generated in this study and the other from the MGI pair-end data generated to estimate the genome size in Kurotani *et al.,* 2023.

| | **Assembly** | | |
|---|---|---|---|
| **Statistics** | **10x** | **10x + ONT** | **PacBio HiFi + Hi-C** |
| **Genome size** | 2,981,676,209 | 2,994,235,606 | 2,926,135,461 |
| **%GC** | 37.96% | 37.96% | 38.12% |
| **Largest contig** | 33,863,123 | 47,606,874 | 184,452,736 |
| **Number of scaffolds** | 58,989 | 50,663 | 1,668 |
| **N50 (L50)** | 5,442,096 (153) | 13,672,338 (68) | 141,754,421 (10) |
| **N90 (L90)** | 22,681 (3,185) | 37,929 (1,810) | 110,643,690 (19) |
| **Completeness (Illumina)** | 96.53% | 97.82% | 99.33% |
| **QV (Illumina)** | 52.64 | 37.88 | 46.43 |
| **Completeness (MGI)** | 96.88% | 96.75% | 98.16% |
| **QV (MGI)** | 46.11 | 37.43 | 43.37 |
| **Complete BUSCOs** | 2257 (97.0%) | 2270 (97.6%) | 2296 (98.7%) |
| **Single copy BUSCOs** | 865 (37.2%) | 859 (36.9%) | 737 (31.7%) |
| **Duplicated BUSCOs** | 1392 (59.8%) | 1411 (60.7%) | 1559 (67%) |
| **Fragmented BUSCOs** | 34 (1.5%) | 28 (1.2%) | 9 (0.4%) |
| **Missing BUSCOs** | 35 (1.5%) | 28 (1.2%) | 21 (0.9%) |

**Table S6. *NbBBLa* genotypes in plants with Cas9-mediated mutations.** The target sequences for each guide RNAs (gRNAs) are provided in the top row; the protospacer adjacent motif (PAM) is underlined, bases in orange text indicate identity to *NbBBLa*. The sequence at the equivalent target locus in each line is shown with mutated bases/deletions in blue. For each line, the location at which the amino acid (aa) sequence first becomes out of frame is given, as well as the position of the first premature stop-codon. Not applicable (NA) indicates that a given gRNA was not included in the construct used to produce this line. Sequences corresponding to mbgRNA17 are not shown as no mutations were detected in any line.

<table>
<tr><th colspan="5">NbBBLa</th></tr>
<tr><th>Plant line</th><th>mbgRNA5<br>TATGAAATCAGAGTAAGGTGCGG</th><th>sgRNA19<br>GTTATTAGACCGAAAAGCCATGG</th><th>sgRNA35<br>GGGAGCTTATATCAACTACTTGG</th><th>mbgRNA97 (rev strand)<br>CCGATTCGCAGCGTCCTCTAAAC</th></tr>
<tr><td>102</td><td>Bi-allelic<br>TATGAAATCAGAGTAA--TGCGG<br>frameshift at aa10, stop codon at aa115<br>TATGAAATCAG-----GGTGCGG<br>frameshift at aa10, stop codon at aa114</td><td>NA</td><td>NA</td><td>NA</td></tr>
<tr><td>138</td><td>NA</td><td>NA</td><td>NA</td><td>Wild type<br>TCGATTTGCTGCATCTTACATGC</td></tr>
<tr><td>159</td><td>NA</td><td colspan="2">Bi-allelic<br>GTTATTAGACCGAAAAG →______ ____________← TACTTGG<br>sequence between sgRNA19 and sgRNA35 is inverted,<br>frameshift at aa214 and stop codon at aa222<br>GTTATTAGACCGAAAAGACCATGG ←75 bp del→<br>frameshift at aa214 and stop codon at aa217</td><td>NA</td></tr>
<tr><td>162</td><td>NA</td><td colspan="2">Homozygous<br>GTTATTAGACCGAAAAGACCATGG ←75 bp del→<br>frameshift at aa214 and stop codon at aa217</td><td>NA</td></tr>
<tr><td>169</td><td>NA</td><td colspan="2">Wild type<br>GTTATTAGACCGAAAAGCCATGG GGGAGCTTATATCAACTACTTGG</td><td>NA</td></tr>
<tr><td>187</td><td>NA</td><td colspan="2">Wild type<br>GTTATTAGACCGAAAAGCCATGG GGGAGCTTATATCAACTACTTGG</td><td>NA</td></tr>
<tr><td>193</td><td>NA</td><td colspan="2">Homozygous<br>GTTATTAGACCGAAAAGACCATGG GGGAGCTTATATCAAC-ACTTGG<br>frameshift at aa214 and stop codon at aa217</td><td>NA</td></tr>
<tr><td>196</td><td>NA</td><td colspan="2">Wild type<br>GTTATTAGACCGAAAAGCCATGG GGGAGCTTATATCAACTACTTGG</td><td>NA</td></tr>
<tr><td>198</td><td>NA</td><td colspan="2">Homozygous<br>GTTATTAGACCGAAAAGACCATGG GGGAGCTTATATCAACTACTTGG<br>frameshift at aa214 and stop codon at aa217</td><td>NA</td></tr>
</table>

**Table S7. *NbBBLb* genotypes in plants with Cas9-mediated mutations.** The target sequences for each guide RNAs (gRNAs) are provided in the top row; the protospacer adjacent motif (PAM) is underlined, bases in orange text indicate identity to *NbBBLb*. The sequence at the equivalent target locus in each line is shown with mutated bases/deletions in blue. For each line and mutation, the location at which the amino acid (aa) sequence becomes out of frame is given, as well as the position of the first premature stop-codon. Not applicable (NA) indicates that a given gRNA was not included in the construct used to produce this line. Sequences corresponding to mbgRNA17 are not shown as no mutations were detected in any line.

| ***NbBBLb*** | | | | |
|---|---|---|---|---|
| **Plant line** | **mbgRNA5**<br>TATGAAATCAGAGTAAGGTGCGG | **sgRNA19**<br>GTTATTAGACCGAAAAGCCATGG | **sgRNA35**<br>GGGAGCTTATATCAACTACTTGG | **mbgRNA97 (rev strand)**<br>CCGATTCGCAGCGTCCTCTAAAC |
| **102** | **Homozygous**<br>TATGAAATCAGAGTAAGGGTGCGG<br>frameshift at aa106 and stop codon at aa120 | NA | NA | NA |
| **138** | NA | NA | NA | **Wild type**<br>tcgatttgctgcatcttacatgc |
| **159** | NA | **Homozygous**<br>GTTATTAGACCGAAAA-CCATGG<br>frameshift at aa218 and stop codon at aa252 | GGGAGCTTATA-------CTTGG | NA |
| **162** | NA | **Homozygous**<br>GTTATTAGACCGAAAA-CCATGG<br>frameshift at aa218 and stop codon at aa252 | GGGAGCTTATA-------CTTGG | NA |
| **169** | NA | **Homozygous**<br>GTTATTAGACCGAAAA-CCATGG<br>frameshift at aa218 and stop codon at aa252 | GGGAGCTTATATCAACTACTTGG | NA |
| **187** | NA | **Wild type**<br>GTTATTAGACCGAAAAGCCATGG | GGGAGCTTATATCAACTACTTGG | NA |
| **193** | NA | **Homozygous**<br>GTTATTAGACCGAAAAGCCATGG | GGGAG------------ACTTGG<br>12 bp in frame deletion replacing 5 residues (AYINY) with a new residue (D) at aa480 | NA |
| **196** | NA | **Homozygous**<br>GTTATTAGACCGAAAAAGCCATGG<br>frameshift at aa218 and stop codon at aa250 | GGGAGCTTATATCAACT---TGG | NA |
| **198** | NA | **Homozygous**<br>GTTATTAGACCGAAAAGACCATGG<br>frameshift at aa218 and stop codon at aa250 | GGGAGCTTATATCAACTACTTGG | NA |

**Table S8. *NbBBLc* genotypes in plants with Cas9-mediated mutations.** The target sequences for each guide RNAs (gRNAs) are provided in the top row; the protospacer adjacent motif (PAM) is underlined and bases in orange text indicate identity to *NbBBLc*. The sequence at the equivalent target locus in each line is shown with mutated bases/deletions in blue. For each line, the location at which the amino acid (aa) sequence first becomes out of frame is given, as well as the position of the first premature stop-codon. Not applicable (NA) indicates that a given gRNA was not included in the construct used to produce this line. Sequences corresponding to mbgRNA17 are not shown as no mutations were detected in any line.

<table>
<tr><th colspan="5">NbBBLc</th></tr>
<tr><th>Plant line</th><th>mbgRNA5<br>TATGAAATCAGAGTAAGGTGCGG</th><th>sgRNA19<br>GTTATTAGACCGAAAAGCCATGG</th><th>sgRNA35<br>GGGAGCTTATATCAACTACTTGG</th><th>mbgRNA97 (rev strand)<br>CCGATTCGCAGCGTCCTCTAAAC</th></tr>
<tr><td>102</td><td>Homozygous<br>TATGAAATCAGAG----GTGCGG<br>frameshift at aa105 and stop codon at aa125</td><td>NA</td><td>NA</td><td>NA</td></tr>
<tr><td>138</td><td>NA</td><td>NA</td><td>NA</td><td>Wild type<br>tcgatttgctgcatcttacatgc</td></tr>
<tr><td rowspan="3">159</td><td rowspan="3">NA</td><td colspan="2">Homozygous</td><td rowspan="3">NA</td></tr>
<tr><td>GTTATTAGACCGA----CCATGG</td><td>GTTATTAGACCGA----CCATGG</td></tr>
<tr><td colspan="2">frameshift at aa218 and stop codon at aa237</td></tr>
<tr><td rowspan="3">162</td><td rowspan="3">NA</td><td colspan="2">Homozygous</td><td rowspan="3">NA</td></tr>
<tr><td>GTTATTAGACCGA----CCATGG</td><td>GTTATTAGACCGA----CCATGG</td></tr>
<tr><td colspan="2">frameshift at aa218 and stop codon at aa237</td></tr>
<tr><td rowspan="2">169</td><td rowspan="2">NA</td><td colspan="2">Wild type</td><td rowspan="2">NA</td></tr>
<tr><td>GTTATTAGACCGAAAAGCCATGG</td><td>GGGAGCTTATATCAACTACTTGG</td></tr>
<tr><td rowspan="3">187</td><td rowspan="3">NA</td><td colspan="2">Homozygous</td><td rowspan="3">NA</td></tr>
<tr><td>GTTATTAGACCGAAAAGCCATGG</td><td>GGGAGCTTATA-------CTTGG</td></tr>
<tr><td colspan="2">frameshift at aa219 and stop codon at aa489</td></tr>
<tr><td rowspan="3">193</td><td rowspan="3">NA</td><td colspan="2">Homozygous</td><td rowspan="3">NA</td></tr>
<tr><td>GTTATTAGACCGAAAA-CCATGG</td><td>GGGAGCTTATA------ACTTGG</td></tr>
<tr><td colspan="2">frameshift at aa219 and stop codon at aa238</td></tr>
<tr><td rowspan="5">196</td><td rowspan="5">NA</td><td colspan="2">Bi-allelic</td><td rowspan="5">NA</td></tr>
<tr><td>GTTATTAGACCGAAAAGCCATGG</td><td>GGGAGCTTATATCAACTTACTTGG</td></tr>
<tr><td colspan="2">frameshift at aa484 and stop codon at aa496</td></tr>
<tr><td>GTTATTAGACCGAAAAGCCATGG</td><td>GGGAGCTTATATCAAC-ACTTGG</td></tr>
<tr><td colspan="2">frameshift at aa484 and stop codon at aa491</td></tr>
<tr><td rowspan="3">198</td><td rowspan="3">NA</td><td colspan="2">Wild type</td><td rowspan="3">NA</td></tr>
<tr><td>GTTATTAGACCGAAAAGCCATGG</td><td>GGGAGCTTATATCAACTACTTGG</td></tr>
<tr><td>n/a</td><td>n/a</td></tr>
</table>

**Table S9. *NbBBLd* genotypes in plants with Cas9-mediated mutations.** The target sequences for each guide RNAs (gRNAs) are provided in the top row; the protospacer adjacent motif (PAM) is underlined and bases in orange text indicate identity to *NbBBLd*. The sequence at the equivalent target locus in each line is shown with mutated bases/deletions in blue. For each line, the location at which the amino acid (aa) sequence first becomes out of frame is given, as well as the position of the first premature stop-codon. Not applicable (NA) indicates that a given gRNA was not included in the construct used to produce this line. Sequences corresponding to mbgRNA17 are not shown as no mutations were detected in any line.

| ***NbBBLd*** | | | | |
|---|---|---|---|---|
| **Plant line** | **mbgRNA5**<br>TATGAAATCAGAGTAAGGTGCGG | **sgRNA19**<br>GTTATTAGACCGAAAAGCCATGG | **sgRNA35**<br>GGGAGCTTATATCAACTACTTGG | **mbgRNA97 (rev strand)**<br>CCGATTCGCAGCGTCCTCTAAAC |
| **102** | **Homozygous**<br>TATGAAATCAGAGTAAG-TGCGG<br>frameshift at aa114 and stop codon at aa235 | NA | NA | NA |
| **138** | NA | NA | NA | **Homozygous**<br>CCGATTACGCAGCGTCCTCTAAAC<br>frameshift at aa77 and stop codon at aa82 |
| **159** | NA | **Wild type**<br>GCTATTAGACCGCAAAGCCATGGG | GGCAGCTTATGTCAACTATATGG | NA |
| **162** | NA | **Wild type**<br>GCTATTAGACCGCAAAGCCATGGG | GGCAGCTTATGTCAACTATATGG | NA |
| **169** | NA | **Wild type**<br>GCTATTAGACCGCAAAGCCATGGG | GGCAGCTTATGTCAACTATATGG | NA |
| **187** | NA | **Wild type**<br>GCTATTAGACCGCAAAGCCATGGG | GGCAGCTTATGTCAACTATATGG | NA |
| **193** | NA | **Wild type**<br>GCTATTAGACCGCAAAGCCATGGG | GGCAGCTTATGTCAACTATATGG | NA |
| **196** | NA | **Wild type**<br>GCTATTAGACCGCAAAGCCATGGG | GGCAGCTTATGTCAACTATATGG | NA |
| **198** | NA | **Wild type**<br>GCTATTAGACCGCAAAGCCATGGG | GGCAGCTTATGTCAACTATATGG | NA |

**Supplementary Table S10. *NbBBLd'* genotypes in plants with Cas9-mediated mutations.** The target sequences for each guide RNAs (gRNAs) are provided in the top row; the protospacer adjacent motif (PAM) is underlined and bases in orange text indicated identity to *NbBBLd'*. The sequence at the equivalent target locus in each line is shown with mutated bases/deletions in blue. For each line, the location at which the amino acid (aa) sequence first becomes out of frame is given, as well as the position of the first premature stop-codon. Not applicable (NA) indicates that a given gRNA was not included in the construct used to produce this line. Sequences corresponding to mbgRNA17 are not shown as no mutations were detected in any line.

<table>
<tr><th colspan="6">NbBBLd'</th></tr>
<tr><th>Plant line</th><th>mbgRNA5<br>TATGAAATCAGAGTAAGGTGCGG</th><th>sgRNA19<br>GTTATTAGACCGAAAAGCCATGG</th><th>sgRNA35<br>GGGAGCTTATATCAACTACTTGG</th><th>mbgRNA 97 (rev strand)<br>CCGATTCGCAGCGTCCTCTAAAC</th></tr>
<tr><td>102</td><td>Homozygous<br>TATGAAATCAGAGTAAG-TGCGG<br>frameshift at aa114 and stop codon at aa238</td><td>NA</td><td>NA</td><td>NA</td></tr>
<tr><td>138</td><td>NA</td><td>NA</td><td>NA</td><td>Wild type<br>CAGATTCGCAGCGTCCTCTAATC</td></tr>
<tr><td>159</td><td>NA</td><td colspan="2">Wild type<br>GCTATTAGACCGCTAAGCCATGGG | GGCAGCTTATGTCAACTATATGG</td><td>NA</td></tr>
<tr><td>162</td><td>NA</td><td colspan="2">Wild type<br>GCTATTAGACCGCTAAGCCATGGG | GGCAGCTTATGTCAACTATATGG</td><td>NA</td></tr>
<tr><td>169</td><td>NA</td><td colspan="2">Wild type<br>GCTATTAGACCGCTAAGCCATGGG | GGCAGCTTATGTCAACTATATGG</td><td>NA</td></tr>
<tr><td>187</td><td>NA</td><td colspan="2">Wild type<br>GCTATTAGACCGCTAAGCCATGGG | GGCAGCTTATGTCAACTATATGG</td><td>NA</td></tr>
<tr><td>193</td><td>NA</td><td colspan="2">Wild type<br>GCTATTAGACCGCTAAGCCATGGG | GGCAGCTTATGTCAACTATATGG</td><td>NA</td></tr>
<tr><td>196</td><td>NA</td><td colspan="2">Wild type<br>GCTATTAGACCGCTAAGCCATGGG | GGCAGCTTATGTCAACTATATGG</td><td>NA</td></tr>
<tr><td>198</td><td>NA</td><td colspan="2">Wild type</td><td>NA</td></tr>
</table>

**Table S11. ANOVA p-vales for Figure 4.** ***, p<0.001; **, p<0.01; *, p<0.05.

| | samples | p-values |
|---|---|---|
| Nicotine uninduced leaves | 138 - 102 | *** |
| | 159 - 102 | |
| | 162 - 102 | |
| | 169 - 102 | *** |
| | 187 - 102 | *** |
| | 193 - 102 | |
| | 196 - 102 | *** |
| | 198 - 102 | |
| | WT - 102 | ** |
| | TC WT - 102 | |
| | CAS9 - 102 | ** |
| | 159 - 138 | *** |
| | 162 - 138 | *** |
| | 169 - 138 | |
| | 169 - 138 | |
| | 187 - 138 | |
| | 193 - 138 | *** |
| | 196 - 138 | |
| | 198 - 138 | *** |
| | WT - 138 | |
| | TC WT - 138 | |
| | CAS9 - 138 | |
| | 162 - 159 | |
| | 169 - 159 | *** |
| | 187 - 159 | ** |
| | 193 - 159 | |
| | 196 - 159 | ** |
| | 198 - 159 | |
| | WT - 159 | * |
| | TC WT - 159 | |
| | CAS9 - 159 | * |
| | 169 - 162 | *** |
| | 187 - 162 | ** |
| | 193 - 162 | |
| | 196 - 162 | ** |
| | 198 - 162 | |
| | WT - 162 | ** |
| | TC WT - 162 | |
| | CAS9 - 162 | * |
| | 187 - 169 | |
| | 193 - 169 | *** |
| | 196 - 169 | |
| | 198 - 169 | ** |
| | WT - 169 | |
| | TC WT - 169 | |

| | | |
|---|---|---|
| | CAS9 - 169 | |
| | 193 - 187 | *** |
| | 196 - 187 | |
| | 198 - 187 | * |
| | WT - 187 | |
| | TC WT - 187 | |
| | CAS9 - 187 | |
| | 196 - 193 | *** |
| | 198 - 193 | |
| | WT - 193 | *** |
| | TC WT - 193 | * |
| | CAS9 - 193 | *** |
| | 198 - 196 | * |
| | WT - 196 | |
| | TC WT - 196 | |
| | CAS9 - 196 | |
| | WT - 198 | |
| | TC WT - 198 | |
| | CAS9 - 198 | |
| | TC WT - WT | |
| | CAS9 - WT | |
| | CAS9 - TC WT | |
| **Nicotine induced leaves** | 138 - 102 | *** |
| | 159 - 102 | |
| | 162 - 102 | |
| | 169 - 102 | *** |
| | 187 - 102 | *** |
| | 193 - 102 | |
| | 196 - 102 | *** |
| | 198 - 102 | |
| | WT - 102 | *** |
| | TC WT - 102 | *** |
| | CAS9 - 102 | *** |
| | 159 - 138 | *** |
| | 162 - 138 | *** |
| | 169 - 138 | |
| | 187 - 138 | |
| | 193 - 138 | *** |
| | 196 - 138 | |
| | 198 - 138 | *** |
| | WT - 138 | * |
| | TC WT - 138 | |
| | CAS9 - 138 | |
| | 162 - 159 | |
| | 169 - 159 | *** |
| | 187 - 159 | *** |
| | 193 - 159 | |
| | 196 - 159 | *** |

| | | |
|---|---|---|
| | 198 - 159 | |
| | WT - 159 | *** |
| | TC WT - 159 | *** |
| | CAS9 - 159 | *** |
| | 169 - 162 | *** |
| | 187 - 162 | *** |
| | 193 - 162 | |
| | 196 - 162 | *** |
| | 198 - 162 | |
| | WT - 162 | *** |
| | TC WT - 162 | *** |
| | CAS9 - 162 | *** |
| | 187 - 169 | |
| | 193 - 169 | *** |
| | 196 - 169 | |
| | 198 - 169 | *** |
| | WT - 169 | ** |
| | TC WT - 169 | ** |
| | CAS9 - 169 | |
| | 193 - 187 | *** |
| | 196 - 187 | |
| | 198 - 187 | *** |
| | WT - 187 | |
| | TC WT - 187 | |
| | CAS9 - 187 | |
| | 196 - 193 | *** |
| | 198 - 193 | |
| | WT - 193 | *** |
| | TC WT - 193 | *** |
| | CAS9 - 193 | *** |
| | 198 - 196 | *** |
| | WT - 196 | |
| | TC WT - 196 | |
| | CAS9 - 196 | |
| | WT - 198 | *** |
| | TC WT - 198 | *** |
| | CAS9 - 198 | *** |
| | TC WT - WT | |
| | CAS9 - WT | |
| | CAS9 - TC WT | |
| **Anabasine uninduced leaves** | 138 - 102 | *** |
| | 159 - 102 | |
| | 162 - 102 | |
| | 169 - 102 | *** |
| | 187 - 102 | ** |
| | 193 - 102 | |
| | 196 - 102 | *** |
| | 198 - 102 | * |

| | | |
|---|---|---|
| | WT - 102 | ** |
| | TC WT - 102 | |
| | CAS9 - 102 | ** |
| | 159 - 138 | *** |
| | 162 - 138 | *** |
| | 169 - 138 | |
| | 187 - 138 | |
| | 193 - 138 | *** |
| | 196 - 138 | |
| | 198 - 138 | |
| | WT - 138 | |
| | TC WT - 138 | |
| | CAS9 - 138 | |
| | 162 - 159 | |
| | 169 - 159 | *** |
| | 187 - 159 | ** |
| | 193 - 159 | |
| | 196 - 159 | *** |
| | 198 - 159 | |
| | WT - 159 | ** |
| | TC WT - 159 | |
| | CAS9 - 159 | ** |
| | 169 - 162 | *** |
| | 187 - 162 | ** |
| | 193 - 162 | |
| | 196 - 162 | *** |
| | 198 - 162 | |
| | WT - 162 | * |
| | TC WT - 162 | |
| | CAS9 - 162 | ** |
| | 187 - 169 | |
| | 193 - 169 | *** |
| | 196 - 169 | |
| | 198 - 169 | |
| | WT - 169 | |
| | TC WT - 169 | |
| | CAS9 - 169 | |
| | 193 - 187 | ** |
| | 196 - 187 | |
| | 198 - 187 | |
| | WT - 187 | |
| | TC WT - 187 | |
| | CAS9 - 187 | |
| | 196 - 193 | *** |
| | 198 - 193 | * |
| | WT - 193 | ** |
| | TC WT - 193 | |
| | CAS9 - 193 | *** |

| | | |
|---|---|---|
| | 198 - 196 | |
| | WT - 196 | |
| | TC WT - 196 | |
| | CAS9 - 196 | |
| | WT - 198 | |
| | TC WT - 198 | |
| | CAS9 - 198 | |
| | TC WT - WT | |
| | CAS9 - WT | |
| | CAS9 - TC WT | |
| **Anabasine induced leaves** | 138 - 102 | *** |
| | 159 - 102 | |
| | 162 - 102 | |
| | 169 - 102 | *** |
| | 187 - 102 | *** |
| | 193 - 102 | |
| | 196 - 102 | *** |
| | 198 - 102 | |
| | WT - 102 | *** |
| | TC WT - 102 | *** |
| | CAS9 - 102 | *** |
| | 159 - 138 | *** |
| | 162 - 138 | *** |
| | 169 - 138 | |
| | 187 - 138 | |
| | 193 - 138 | *** |
| | 196 - 138 | |
| | 198 - 138 | *** |
| | WT - 138 | |
| | TC WT - 138 | |
| | CAS9 - 138 | |
| | 162 - 159 | |
| | 169 - 159 | *** |
| | 187 - 159 | *** |
| | 193 - 159 | |
| | 196 - 159 | *** |
| | 198 - 159 | |
| | WT - 159 | *** |
| | TC WT - 159 | *** |
| | CAS9 - 159 | *** |
| | 169 - 162 | *** |
| | 187 - 162 | *** |
| | 193 - 162 | |
| | 196 - 162 | *** |
| | 198 - 162 | |
| | WT - 162 | *** |
| | TC WT - 162 | *** |
| | CAS9 - 162 | *** |

| | | |
|---|---|---|
| | 187 - 169 | |
| | 193 - 169 | *** |
| | 196 - 169 | |
| | 198 - 169 | *** |
| | WT - 169 | |
| | TC WT - 169 | |
| | CAS9 - 169 | |
| | 193 - 187 | *** |
| | 196 - 187 | |
| | 198 - 187 | *** |
| | WT - 187 | |
| | TC WT - 187 | |
| | CAS9 - 187 | |
| | 196 - 193 | *** |
| | 198 - 193 | |
| | WT - 193 | *** |
| | TC WT - 193 | *** |
| | CAS9 - 193 | *** |
| | 198 - 196 | *** |
| | WT - 196 | |
| | TC WT - 196 | |
| | CAS9 - 196 | |
| | WT - 198 | * |
| | TC WT - 198 | |
| | CAS9 - 198 | * |
| | TC WT - WT | |
| | CAS9 - WT | |
| | CAS9 - TC WT | |
| **Anatabine uninduced leaves** | 169 - 138 | |
| | 187 - 138 | |
| | 196 - 138 | |
| | 198 - 138 | |
| | WT - 138 | |
| | TC WT - 138 | ** |
| | CAS9 - 138 | |
| | 187 - 169 | |
| | 196 - 169 | |
| | 198 - 169 | |
| | WT - 169 | |
| | TC WT - 169 | |
| | CAS9 - 169 | |
| | 196 - 187 | |
| | 198 - 187 | |
| | WT - 187 | |
| | TC WT - 187 | |
| | CAS9 - 187 | |
| | 198 - 196 | |
| | WT - 196 | |

| | | |
|---|---|---|
| | TC WT - 196 | |
| | CAS9 - 196 | |
| | WT - 198 | |
| | TC WT - 198 | |
| | CAS9 - 198 | |
| | TC WT - WT | |
| | CAS9 - WT | |
| | CAS9 - TC WT | |
| **Anatabine induced leaves** | 169 - 138 | |
| | 187 - 138 | |
| | 196 - 138 | |
| | 198 - 138 | *** |
| | WT - 138 | |
| | TC WT - 138 | ** |
| | CAS9 - 138 | |
| | 187 - 169 | |
| | 196 - 169 | |
| | 198 - 169 | *** |
| | WT - 169 | |
| | TC WT - 169 | * |
| | CAS9 - 169 | |
| | 196 - 187 | |
| | 198 - 187 | *** |
| | WT - 187 | |
| | TC WT - 187 | * |
| | CAS9 - 187 | |
| | 198 - 196 | *** |
| | WT - 196 | |
| | TC WT - 196 | * |
| | CAS9 - 196 | |
| | WT - 198 | *** |
| | TC WT - 198 | *** |
| | CAS9 - 198 | *** |
| | TC WT - WT | |
| | CAS9 - WT | |
| | CAS9 - TC WT | |

**Table S12. ANOVA p-values for Figure 6.** ***, p<0.001; **, p<0.01; *, p<0.05.

| | samples | p-values |
|---|---|---|
| **nicotine** | Cas9 - 102 | *** |
| | WT - 102 | *** |
| | WT - Cas9 | |
| **anabasine** | Cas9 - 102 | *** |
| | WT - 102 | *** |
| | WT - Cas9 | * |
| **anatabine** | Cas9 - 102 | *** |
| | WT - 102 | *** |
| | WT - Cas9 | |

**Figure S1. Schematic and photographs of the hydroponic system.** For precursor feeding experiments and root harvesting, *N. benthamiana* seedlings were germinated and grown on cotton gauze (28 thread) glued onto rubber O-rings (¾ inch or 1 ¼ inch) fitted onto the wells of multi-well plates (6 or 12 wells, respectively).

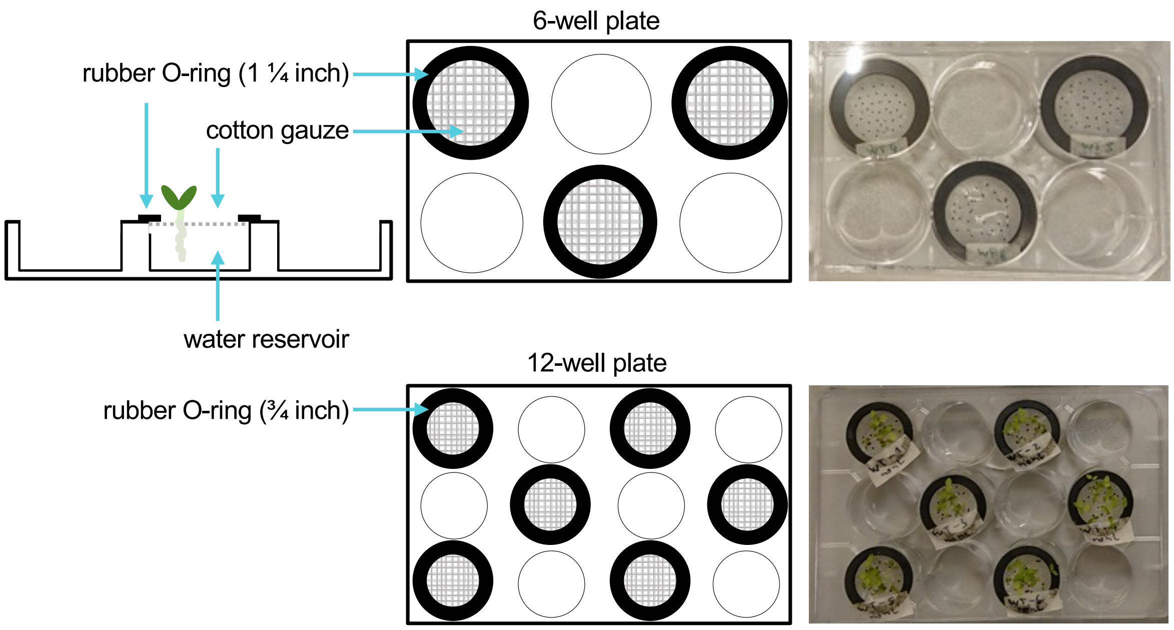

**Figure S2. Maximum likelihood tree and multiple sequence alignment of *N. tabacum* and *N. benthamiana* BBLs.** The phylogeny was reconstructed using PhyML3.0. The premature stop codon in NbBBLd' is indicated by red, capitalized text.

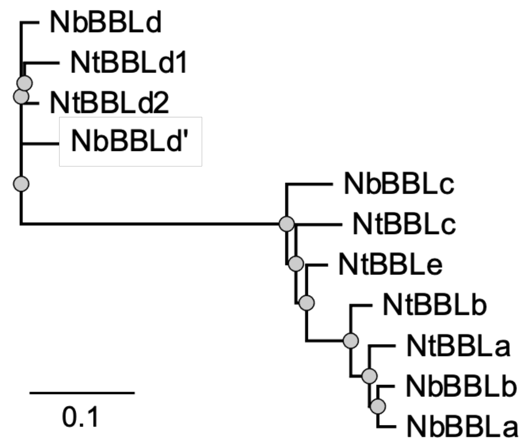

```
Consensus atg---------------------------tttccgctcataattct------gatcagc    27
NbBBLc    atg---------------------------tttctactcataattct------gatcagc    27
NbBBLa    atg---------------------------tttccgctcataattct------catcagc    27
NbBBLb    atg---------------------------tttccactcataattct------gatcagc    27
NtBBLc    atg---------------------------tttccactcataattct------aatcagc    27
NtBBLe    atg---------------------------tttccaatcataattct------gatcagc    27
NtBBLa    atg---------------------------tttccgctcataattct------gatcagc    27
NtBBLb    atg---------------------------tttccactcataattct------gatcagc    27
NbBBLd    atgaaacgaaatatatccatgtttcttcagcttctgctcataattctgatgatgatcagc    60
NbBBLd'   atgaagcgaaatatatccatgtttcttcagcttctgctcataattctaatgatgatcagc    60
NtBBLd1   atgaaacgaaatatatccatgtttcttcagcttctgctcattattctgatgatgatcagc    60
NtBBLd2   atgaaacgaaatatatccatgtctcttcagcgtttgctcataattctgatgatgatcagc    60

Consensus ttttcatttacttccctctctgctactgcta------ctagtggagca---g---gagga    75
NbBBLc    ttttcacttacttcccgctctgctactgctactgctactagtggagca---g---aagga    81
NbBBLa    ttttccc---------tctctgctactgcta------ctagtggagca---g---gagga    66
NbBBLb    ttttcacttacttccctctctgctactgcta------ctagtggagca---gaaggagga    78
NtBBLc    ttctcatttactttcctctctgctagtgcta------ctagtggagca---ggagaagga    78
NtBBLe    ttttcatttactttcctctttgctagtgtta------ctagtggagca---g---gagga    75
NtBBLa    ttttcacttgcttccttgtctgaaactgcta------ct------------g---gagct    66
NtBBLb    ttttcacttacttccctctctgctactgcaa------ctagtggagcaggag---gtgga    78
NbBBLd    ttcttatttacttctcttcttgtaccatccg------tctct---gca---a---caaat   105
NbBBLd'   ttcttatttacttgtcttcttgtatcttcca------tctct---gca---a---caaat   105
NtBBLd1   ttcttatttacttctcttcttgtaccttcgg------tctct---gca---a---caact   105
NtBBLd2   ttcttatttacttctcttcttgtaccttccg------tctct---gct---a---caaat   105

Consensus gttacaaatctttccacctgtttaatcaaccacaatgtccataacttctctatttacccc   135
NbBBLc    gttacaaatctttccacctgtttaatcaaccacaatgtccataacttctctatataccccc   141
NbBBLa    gttacaaatctttcaacctgtttaatcaaccacaatgtccataacttctctatttacccc   126
NbBBLb    gttacaaatctttcaacctgtttaaccaaccacaatgtctataacttctctatttacccc   138
NtBBLc    gttgcaaatctttccacctgtttaatcaaccacaatgtccataatttctccatgtacccc   138
NtBBLe    gttacaaatctttccacctgtttaatcaaccacaatgtccataacttctctatttacccc   135
NtBBLa    gttacaaatctttcagcctgcttaatcaaccacaatgtccataacttctctatttacccc   126
NtBBLb    gttgcaaatctttacacctgtttaatcgaccacaatgtccataacttctctatttacccc   138
NbBBLd    ctcaataccatttccacctgtttattcaattataaagtcactaacttctctgtttaccca   165
NbBBLd'   ctcaataccatttccacctgtttaatcaattacaaagtcagtaacttttctgtttaccca   165
NtBBLd1   ctcaataccatttccacctgtttaatcaattacaaagtcagtaacttctctgtttaccca   165
NtBBLd2   ctcaataccatttccacctgtttgatcaattacaaagtcagtaacttctctgtttatcca   165

Consensus acaaggaatgatccaa---gtagtaattactttaacttgctcgatttctcccttcagaat   192
NbBBLc    acaaagaatgatccaa---gtagaaattactttaacttgctcgatttctcccttcaaaat   198
NbBBLa    acaaggaacgatccaa---atagtaattactttaacttgctcgacttctcccttcagaat   183
NbBBLb    agaaggaatgatccaa---atagttcttactttaacttgctcgacttctcccttcagaat   195
NtBBLc    acaag---------------tagaaattactttaacttgctcgacttctcccttcagaat   183
NtBBLe    acaaagaatgatcaaagtagtagtaattactttaacttgctcgatttttcccttcagaat   195
NtBBLa    acaag---------------tagaaattactttaacttgctccacttctcccttcaaaat   171
NtBBLb    acaaagaatgatcaaagtagtagtaattactttaacttgctcgatttttcccttcagaat   198
NbBBLd    acaaggaatcatgctg---gtaatagttactataacttgcttgatttctccattcagaat   222
NbBBLd'   acaaggaatcatgctg---ataatagttactacaacttgcttgatttctccattcagaat   222
NtBBLd1   acaaggaatcatgctg---gtaatagttactataacttgcttgatttctccattcagaat   222
NtBBLd2   acaaggaatcatgctg---gtaataggtactataacttgcttgatttctccattcagaat   222

Consensus cttcgatttgcagcatcttacatgccgaaaccaacggtcattatcctaccaaacagcaag   252
NbBBLc    cttcgatttgctgcatcttacatgccgaaaccaacggtcattatcctaccaagcagcaag   258
```

| | | |
|---|---|---|
| NbBBLa | cttcgatttgctgcatcttacatgccgaaaccaaccttcattatcctaccaagcagcaag | 243 |
| NbBBLb | cttcgatttgcagcatcttacatgccgaaaccaacgttcattatcctaccaagcagcaag | 255 |
| NtBBLc | cttcgatttgcagcatctaacatgccgaaaccaacggtcattatcctaccaaacagcaag | 243 |
| NtBBLe | cttcgatttgctgcatcttacatgccgaaaccaacggtcattatcctaccaaacagcaaa | 255 |
| NtBBLa | cttcgctttgctgcacctttcatgccgaaaccaaccttcattatcctaccaagcagtaag | 231 |
| NtBBLb | cttcgatttgctgcatcttacatgccgaaaccaacggtcattattctaccaaacagcaaa | 258 |
| NbBBLd | ctccgattcgcagcgtcctctaaaccaaaaccaacggtcattattgtaccagagagcaag | 282 |
| NbBBLd' | ctcagattcgcagcgtcctctaatccaaagccaactgttattattgtaccagagatcaag | 282 |
| NtBBLd1 | ctccgattcgcagcgtgctctaaaccaaaaccaactgtcattatcgtaccagagagcaag | 282 |
| NtBBLd2 | ctccgattcgcagcgtcctctaaaccaaaaccaacggtcattatcgtaccagagagcaag | 282 |
| Consensus | gaggagctcgtgagcaccattctttgttgcagacaagcatcttatgaaatcagagtaagg | 312 |
| NbBBLc | gaggagctcgtgagcaccatttctgttgcagaaaagcatcttatgaaatcagagtaagg | 318 |
| NbBBLa | gaggagctcgtgagcaccatttttgttgcagaaaagcatcttatgaaatcagagtaagg | 303 |
| NbBBLb | gaggagctcgtgagcaccatttttgttgcagaaaagcatcttatgaaatcagagtaagg | 315 |
| NtBBLc | gaggagctcgtgagcaccattctttgttgcagacaaacatcttatgaaatcagagtaagg | 303 |
| NtBBLe | gaggagctcgtgagtaccattctttgttgcagacaaacatcttatgaaatcagagtaagg | 315 |
| NtBBLa | gaggagctcgtgagcaccatttttgttgcagaaaagcatcttatgaaatcagagtaagg | 291 |
| NtBBLb | gaggagctcgtgagcaccattctttgttgcagacaagcttcttatgaaatcagagtaagg | 318 |
| NbBBLd | gagcagctggtgagcagcgttctgtgttgcagacaaggttcttatgaaatcagagtaagg | 342 |
| NbBBLd' | gagcagctggtgagcagcgttctgtgttgcagacaaggttcttatgaaatcagagtaagg | 342 |
| NtBBLd1 | gagcagctggtgagcagcgttctgtgttgcagacaaggctcgtatgaaatcagagtaagg | 342 |
| NtBBLd2 | gagcagctggtgagcagcgttctgtgttgcagacaaggttcttatgaaatcagagtaagg | 342 |
| Consensus | tgcggaggacacagttacgaaggaacttcttacgtttcctttgacggttccccattcgtg | 372 |
| NbBBLc | tgcggaggacacagttatgaaggaacttcttacgtttcctttgacggttctcaattcgtg | 378 |
| NbBBLa | tgcggcggacacagttacgagggaacatcttacgtttcctttgatgcttccccttcgtg | 363 |
| NbBBLb | tgcggcggacacagttacgagggaacatcttacgtttcctttgacggttccccgttcgtg | 375 |
| NtBBLc | tgcggaggacacagttacgaaggaacttcttctgtttcctttgacggttccccgttcgtg | 363 |
| NtBBLe | tgcggaggacacagttacgagggaacttcttacgtttcctttgacggttcccctttcgtg | 375 |
| NtBBLa | tgcggcggacacagttacgaaggaacttcttacgtttcctttgacgcttctccattcgtg | 351 |
| NtBBLb | tgcggaggacatagttacgagggaacttcttacgtttcctttgacggttccccgttcgtg | 378 |
| NbBBLd | tgcggaggacacagttatgaaggaacttcttacgtttcctttgatggttccccatttgtg | 402 |
| NbBBLd' | tgcggaggacacagttatgaagggacttcatacgtttcctttgatggttccccatttgtt | 402 |
| NtBBLd1 | tgcggtggacacagttatgaagggacttcatcagtttcctttgatggttccccatttgtg | 402 |
| NtBBLd2 | tgcggaggacacagttatgaagggacttcttacgtttcctttgatggttccccatttgtg | 402 |
| Consensus | atcgttgacttgatgaaattagacgacgtttcagtagatttggattccgaaacagcttgg | 432 |
| NbBBLc | atcgttgacttgatgaaattagacgacgtttcagtagatttggattccgaaacagcttgg | 438 |
| NbBBLa | atcgttgacttgatgaaattagacaacgtttcagtagatttggattccgaaactgcttgg | 423 |
| NbBBLb | atcgttgacttgatgaaattagacgacgtttcagtagatttggattctgaaactgcttgg | 435 |
| NtBBLc | atcatcgacttgatgaaattagacgacgtttcagtagatttggattccgaaactgcttgg | 423 |
| NtBBLe | atcgttgacttgatgaaattagacgacgtttcagtagatttggattccgaaacagcttgg | 435 |
| NtBBLa | atcgttgacttgatgaaattagacgacgtttcagtagatttggattctgaaacagcttgg | 411 |
| NtBBLb | atcgttgacttgatgaaattagacgaagtttcagtagatttggattctgaaactgcttgg | 438 |
| NbBBLd | gtaattgatttaatgaaattagacggcatttcagtagatttggattccgaaacagcttgg | 462 |
| NbBBLd' | gtcattgatttgatgaaattagacggcgtttcagtagatttggattccgaaacagca**TGA** | 462 |
| NtBBLd1 | gtcattgatttgatgaaattagacggcgtttcagtggatgtggattcagaaaccgcgtgg | 462 |
| NtBBLd2 | gtcattgatttgatgaaattagatgatgtttcggtagatttggattccgaaaccgcgtgg | 462 |
| Consensus | gctcagggcggcgcaacaattggccaaatttattacgccattgccaaggtaagtgacgtt | 492 |
| NbBBLc | gcccagggcggcgcaacaattggccaaatttattacgccattgccaaggtaagtgatgtt | 498 |
| NbBBLa | gcccagggcggcgcaacaattggccaaatttattacgccattgccaaggtaagtgacgtt | 483 |
| NbBBLb | gctcagggaggcgcaacaattggccaaatttattacgccattgccaaggcaagtgacgtt | 495 |
| NtBBLc | gctcagggcggcgcaacaattggccaaatttattacgccattgccaaggcaagtgacgtt | 483 |
| NtBBLe | gctcagggcggcgcaacaattggccaaatttattacgccatttccagggttagtgacgtt | 495 |
| NtBBLa | gctcagggcggcgcaacaattggccaaatttattatgccattgccaaggtaagtgacgtt | 471 |
| NtBBLb | gctcagggcggcgcaacaattggccaaatttattacgccattgccaaggtaagtgacgtt | 498 |
| NbBBLd | gtacaaggtggcgctacacttggccagacttattatgccatttcccaggccagcgacgtt | 522 |
| NbBBLd' | gtacaaggtggcactacacttggccagacttattatgccatttcccaggtcagtggcgtt | 522 |
| NtBBLd1 | gtacagggcggcgctacacttggccagacttattatgccatttcccgagccagcaacgtt | 522 |
| NtBBLd2 | gtacaaggtggcgctacacttggccagacttattatgccatttcccgggccagtgacgtt | 522 |
| Consensus | catgcattttcagcaggttcgggaccaacagtaggatctggaggtcatatttcaggtggc | 552 |
| NbBBLc | catgcattttcagcaggttcggcatcaacagtaggatctggcggtcatatttcaggtggc | 558 |
| NbBBLa | catgcattttcagcaggttcgggaccaaccgtaggatctggaggtcatatttccggtggc | 543 |
| NbBBLb | catgcattttcagcgggttcggggccaacagtaggatctggaggtcatatttccggtggc | 555 |
| NtBBLc | catgcattttcagcaggttcgggtccaacagtaggatctggaggtcatatttcaggtggc | 543 |
| NtBBLe | catgcattttcagcaggttcgggaccaacagtaggatctggaggtcatatttcaggtggc | 555 |
| NtBBLa | catgcattttcagcaggttcgggaccaacagtaggatctggaggtcatatttcaggtggt | 531 |
| NtBBLb | catgcattttcagcaggttcgggaccaacagtaggatctggaggtcatatttcaggtggc | 558 |
| NbBBLd | catggattttcagctggttcttgcccaacagttggggttgggggccacatttccgggggt | 582 |
| NbBBLd' | catggattttcagctggttcttgcccaacagttggggtcgggggccacatttcagggggt | 582 |
| NtBBLd1 | catggattttcagctggttcttgcccaacagttggggttggcgggcacatttccgggggt | 582 |
| NtBBLd2 | catggattttcagctggttcttgcccaacagttggggttgggggccacatttccgggggt | 582 |
| Consensus | ggctttggacttttatcyagaaaattcggacttgctgctgataatgtcgttgatgctctt | 612 |
| NbBBLc | ggctttggacttttgtccagaaaattcggactcgctgcggataatgttgttgatgctctt | 618 |
| NbBBLa | ggatttggacttttatctagaaaattcggacttgctgctgataatgtcgttgatgctctt | 603 |
| NbBBLb | ggatttggacttttatctaggaaattcggacttgctgctgataatgtcgttgatgctctt | 615 |

```
NtBBLc    ggctttggacttctgtccagaaaattcggagtcgctgctgatagtgtcgttgatgctctt   603
NtBBLe    ggctttggactaatgtccagaaaattcggactcgctgctgatagtgtcgttgatgctctt   615
NtBBLa    ggatttggacttttatctagaaaattcggacttgctgctgataatgtcgttgatgctctt   591
NtBBLb    ggctttggacttttatctagaaaattcggactcgctgctgataatgtcgttgatgctctt   618
NbBBLd    ggctttggatttttatcaagaaaatatggacttgctgctgataacgtggtcgatgctctt   642
NbBBLd'   ggctttggatttttgtcaaggaaatatggacttgctgctgataacgtggtggatgctctt   642
NtBBLd1   ggttacggatttttatccagaaaatatggacttgctgctgataacgtggtggatgctctt   642
NtBBLd2   ggctttggatttttatcaagaaaatatggacttgctgctgataacgtggtggatgctctt   642

Consensus cttattgatgctgaaggacggttattagaccgaaaagccatgggagaagacgtgttttgg   672
NbBBLc    cttattgatgctgaaggacggttattagaccgaaaagccatgggagaggacgtattttgg   678
NbBBLa    cttattgatgctgatggacggttattagaccgaaaagccatgggtgaagacgtgttttgg   663
NbBBLb    ctaattgatgctgatggacggttattagaccgaaaagccatgggagaagacgtgttttgg   675
NtBBLc    cttattgatgctgatggacggttattagaccgaaaagccatgggagaagacgtgttttgg   663
NtBBLe    ctaattgatgctgaaggacggttattagaccggaaagccatgggagaagacgtattttgg   675
NtBBLa    cttattgatgctgatggacggttattagaccgaaaagccatgggcgaagacgtgttttgg   651
NtBBLb    cttatcgatgctgatgggcggttattagaccgaaaagccatgggagaagacgtgttttgg   678
NbBBLd    cttgttgattcggaaggacggctattagaccgcaaagccatgggagaagaagtgttttgg   702
NbBBLd'   cttgttgatgcggaaggacggctattagaccgctaagccatgggagaagaagtgttttgg   702
NtBBLd1   cttgttgatgcggaaggacggctattagaccgcaaagccatgggagaagaaatcttttgg   702
NtBBLd2   cttgttgatgcggaaggacggctattagaccgcaaagccatgggagaagaagtgttttgg   702

Consensus gcaatcagaggtggcggyggtggaaattggggaattatttatgcctggaaaattcgatta   732
NbBBLc    gcaatcagaggtggaggcggtggaaattggggaataatttatgcctggaaaatcagatta   738
NbBBLa    gcaatcagaggtggcggtggtggaaattggggaattgtttatgcctggaaaattcgatta   723
NbBBLb    gcaatcagaggtggcggcggtggaaattggggaattgtttatgcctggaaaattcgatta   735
NtBBLc    gcaatcagaggtggcggcggtggaaattggggaattatttatgcctggaaaattcgatta   723
NtBBLe    gcaatcagaggtggcggcggtggaaattggggaattatttatgcctggaaaattcgatta   735
NtBBLa    gcaatcagaggtggcggcggtggaaattggggcattgtttatgcctggaaaattcgatta   711
NtBBLb    gcaatcagaggtggcggaggaggaaattggggaattatttatgcctggaaaattcgatta   738
NbBBLd    gccatcagaggtggtggtggaggaatttggggaatcatttatgcctggaaaatccgattg   762
NbBBLd'   gccatcagaggtggtggtggagggatttggggaatcatttacgcctggaaaatccgattg   762
NtBBLd1   gccatcagaggtggaggtggaggaatttggggaatcatttacgcctggaaaatccgattg   762
NtBBLd2   gccatcagaggtggtggtggaggaatttggggaatcatttacgcctggaaaatccgattg   762

Consensus ctcaaagtgcctaaaatcgtaacaacttttatgatctctaggcctggmtccaaacaatac   792
NbBBLc    atcaaagtgcctaaaatcgtaacaacttttgtgatctctaggcctggctccaaacaatac   798
NbBBLa    ctcaaagtgcctaaaattgtaacagcttgtatgatctataggcctggatccaaacaatac   783
NbBBLb    ctcagagtgcctaaaatcgtaacagcttgtatgatctataggcctggatccaaacaatac   795
NtBBLc    gtgaaagtgcccaaaatcgtaacaacttttaagatctctaagcctggctccaaacaatac   783
NtBBLe    ctcaaagtgcctaaaatcgtaacaacttgtatgatctataggcctggatccaaacaatac   795
NtBBLa    ctcaaagtgcctaaaatcgtaacaacttgtatgatctataggcctggatccaaacaatac   771
NtBBLb    ctcaaagtgcctaaaatcgtaacaacttgtatgatctataggcctggatccaaacaatac   798
NbBBLd    ctcaaagtgcccaagaccgtgactagtttcatagtccctaggcctggctccagacgatat   822
NbBBLd'   ctcaaagtgcccaagactgtgactagtttcataatccctaggcctgggtccaaacgat--   820
NtBBLd1   ctcaaagtgcccaagaccgtgaccagtttcataatccctaggcctggctccaaacgatat   822
NtBBLd2   ctcaaagtgcccaagactgtgactagtttcatagtccctaggcctggctccaaacgatat   822

Consensus gtggcccaactacttcacaaatggcaaatagttgcaccaaatttggacgatgattttact   852
NbBBLc    gttgcctcattacttcacaaatggcaattagttgcaccaaatttggacgatgattttact   858
NbBBLa    gtggctcaaatacttgagaaatggcaagtagttactccaaatttagtcgatgattttact   843
NbBBLb    gtggctcaaatacttgagaaatggcaaattgttactccaaatttggtcgatgattttact   855
NtBBLc    gttgccccattactttacaaatggcaaatagttgcaccaaatttggccgatgattttact   843
NtBBLe    gtggctcaactacttcagaaatggcaaatagttactccaaatttggccgatgattttact   855
NtBBLa    gtggctcaaatacttgagaaatggcaaatagttactccaaatttggtcgatgattttact   831
NtBBLb    gtggctcaactacttcagaaatggcaaatagttactccaaatttggtcgatgattttact   858
NbBBLd    gtgtcccaactagttcacaaatggcaacttgttgcaccaaagttagacgatggcttttat   882
NbBBLd'   gtgtcccaactagttcacaaatggcaacttgttgcaccaaagttagacgatgacttttat   880
NtBBLd1   gtgtcccaactagttcacaaatggcaacttgttgcaccaaagttagaggatgaattttat   882
NtBBLd2   gtgtcccaactagttcacaaatggcaacttgttgcaccaaagttagacgatgacttttat   882

Consensus ctaggagtactcatgagacctgcaaatctnccggcggatatgaaatawgg-aatamtact   911
NbBBLc    ctaggagtagggatgggaccattcaatctgccagcggatatgaaatacggaaatactact   918
NbBBLa    ctaggagtactcctgaggcctgcagatctaccggcggatatgaaatatggtaatagtact   903
NbBBLb    ctaggagtactcctgagacctgcaaatctaccggcggatatgaaatatggtaatagtact   915
NtBBLc    ctaggagtacaaatgatacctatagatctgccggctgatatgaaatacggaaatcctact   903
NtBBLe    ctaggagtactcatgagacctatagatctgcgggcggatatgaattacggaaatactact   915
NtBBLa    ctaggagtactgctgagacctgcagatctacccgcggatatgaaatatggtaatactact   891
NtBBLb    ctaggagtactcttgagacctgcagatctaccggcggatatgaaatatggcaacagtact   918
NbBBLd    ctatcgatctccatgagctctgctagtaa------------------agg---aaacatt   921
NbBBLd'   atatcgatctccatgagctctgctagtaa------------------agg---aaacatt   919
NtBBLd1   ctatcgatctccatgagctctcctagtaa------------------agg---aaacatt   921
NtBBLd2   ctatcgatctccatgagctctgctagtaa------------------agg---aaacatt   921

Consensus cctattgaaatatttccccaattcaacgcactttatttgggtccaaaaactgaagccatt   971
NbBBLc    cctgttgaaataattccccaattcaacgcactttatttgggtccaaaaactgaagctatt   978
NbBBLa    cctattgaaatatttccccaattcaacgcactttatttgggtccaaaaactgaagttctt   963
NbBBLb    cctattgaaatatttccccaattcaacgcgctttatttgggtccaaaaactgaagctctt   975
NtBBLc    cctattgaaatatgtccccaattcaatggactttatctgggtccaaaaactgaagcggtt   963
NtBBLe    cctattgaaacatttccccaattcaatgcactttatttgggtccaaaaactgaagcggtt   975
```

```
NtBBLa    cctattgaaatatttccccaattcaatgcactttatttgggtccaaaaactgaagttctt   951
NtBBLb    cctattgaaatatttccccaattcaatgcactttatttgggtccaaaaactgaagtcctt   978
NbBBLd    cctattgaagtaaatgcccaattcagcggattttaccttggtacaaaaaccgaagccatt   981
NbBBLd'   ccttttgaaataaatgcccaattcagcggattttacttaggtacaaaaaccgaagccatt   979
NtBBLd1   cctattgaaataaatgcccaattcagcggattttacctaggtacaaaaaccgaagccatt   981
NtBBLd2   cctattgaaataaatgcccaattcagcggattttacctaggtacaaaaaccgaagccatt   981

Consensus tccatattgaatgaggcatttccggagctgggcgttaagaatgatgactgcaaagaaatg  1031
NbBBLc    tccatattaaatgaggcatttccagagctggacgttaagaatgatgacggcaaagaaatg  1038
NbBBLa    tccatatcaaatgagacatttccggagctaggcgttaagagtgatgagtgcaaggaaatg  1023
NbBBLb    tccatatcaaatgagacatttccggagctaggcgttaagaatgacgagtgcacggaaatg  1035
NtBBLc    tctatattaaatgaggcatttccagagctgaacgttaagaatgatgacgccaaagaaatg  1023
NtBBLe    tccatattaaatgaggcatttccagagctggacgctaagaatgatgacgccaaagaaatg  1035
NtBBLa    tccatatcgaatgagacatttccggagctaggcgttaagaatgatgagtgcaaggaaatg  1011
NtBBLb    tctatatcgaatgaggaatttccggagctgggcgttaagaatgatgagtgcaaggaaatg  1038
NbBBLd    tccatcttgaatgaggcctttccggaggtcggagttctggaagatgactgcaaagaaatg  1041
NbBBLd'   tccatcttgaacgaggcctttccggaggtgggagttctggaagatgactgcatagaaatg  1039
NtBBLd1   tccatcttgaatgaggccttttcggagttgggagttctggaaggtgactgcaaagaaatg  1041
NtBBLd2   tccatcttgaatgaggcctttccggagttgggagttgtggaaagtgactgcaaagaaatg  1041

Consensus acttggattgagtcagcacttttcttctccgaattagataacgtta-cgggaactcctct  1090
NbBBLc    acttggatagagtcagcacttatcttttccgaatcagctaacgtattcgggaactcttcc  1098
NbBBLa    acttgggtagagtcatcactttacttctccgaattagctgacgttagcgggaactcctct  1083
NbBBLb    acttgggtagagtcatcactttacttctccgaattagctgacgttagcgggaactcctct  1095
NtBBLc    acttggattgagtctgcacttttcttttccgacttagataacatattcgggaactcctct  1083
NtBBLe    acttggattgagtcagcacttttcttttccgaattagataacgtattcgggaactcctct  1095
NtBBLa    acttgggtagagtcagcacttttcttctccgaattagctgacgttaacgggaactcgact  1071
NtBBLb    acttggatagagtcagcacttttcttctccgaattagctgacattaacgggaattcctct  1098
NbBBLd    agttggattgaatcaacacttttcttctccgaattagataacgttg---cgaacacctc-  1097
NbBBLd'   agttggattgaatcaacgcttttcttctccaaattagataacggta---cgaacacctc-  1095
NtBBLd1   agttggattgaatcaacacttttcttctccgaattaaatgacgttg---cgaattcctc-  1097
NtBBLd2   agttggattgaatcaacacttttcttctccgaattagataacgttg---cgaacacctc-  1097

Consensus gncgatatctcccgtttgaaagaacgttacatggacggaaaatctttcttcaaaggcaaa  1150
NbBBLc    gacgatatctcccgtttgagagaacgctacacggacggaaaatctttcttcaaatgcaaa  1158
NbBBLa    ggtgatatctcccgtctgaaagaacgttacatggacggaaaaggtttcttcaaaggcaag  1143
NbBBLb    gctgatatctcccgtctgaaagaacgttacatggacggaaaaggtttcttcaaaggcaag  1155
NtBBLc    gacgatatctcccatttgaaagaacgctacttgggtgtaaaaatttgcttcaaaggcaaa  1143
NtBBLe    gacgatatctcccgtttgaaagaacgctacatggacgcaaaaactttcttcaaaggcaaa  1155
NtBBLa    ggtgatatctcccgtctgaaagaacgttacatggacggaaaaggtttttttcaaaggcaaa  1131
NtBBLb    aatgatatctcccgtctgaaagaacgttacatggacggaaaaggtttcttcaaaggcaaa  1158
NbBBLd    --cgatgtctctcgtctaaaagaacgttactttgaaaacaaatcatacttcaaagccaaa  1155
NbBBLd'   --cgatgtctctcgtttgaaagagcgttactttgaaaaccaatcatacttcaaagccaaa  1153
NtBBLd1   --cgatgtctctcgtttgaaagagcgttactttgaaaacaaatcatacttcaaagccaaa  1155
NtBBLd2   --cgatgtctctcgtttgaaagagcgttactttgaaaacaaatcatacttcaaagccaaa  1155

Consensus tcagactatgtgaagaccccartttcaatggatgggatgatgacagctcttgttgaactc  1210
NbBBLc    tcagattatgtgaagatcccattttcaatggacgggatgatgacagctcttgttgaacta  1218
NbBBLa    acggactatgtgaagaagccagtttcaatggatgggatgctcacatttcttgttgaactc  1203
NbBBLb    acggactatgtgaagaagccagtttcaatggatgggatgctcacatttcttgttgaactc  1215
NtBBLc    tcagattatgtgaagaccccattttcaatggacggaataatgacagctcttgttgaacac  1203
NtBBLe    tcagattttgtgaagactccattttcaatggacgcgatgatgacagctcttgttgaactc  1215
NtBBLa    acggactacgtgaagaagccagtttcaatggatgggatgctaacatttcttgtggaactc  1191
NtBBLb    acggactatgtgaagaagccagtttcaatggatgggatgctaacatttcttgttgaactc  1218
NbBBLd    tcagactatgtgaagaccccaatttcagtgggagggattatgacagctcttgatgttctt  1215
NbBBLd'   tcagactatgtgaagaccccaatttcagtgggagggattatgacagctcttgatgttctt  1213
NtBBLd1   tcagactatgtgaagaccccaatttcagtgggtgggattatgacggctcttaatgttctt  1215
NtBBLd2   tcagaccatgtgaagaccccaatttcagtgggagggattatgacagctcttgatgttctt  1215

Consensus gagaaaaaccccaagggatatcttgtcttygatccttatggcggagccatggacaagatt  1270
NbBBLc    gagaaaaaacccaactcataccttatcttcgatccttatggtggagccatggacaagatt  1278
NbBBLa    gagaaaaacccgaagggataccttgtcttcgatccttatggcggagccatggacaagatt  1263
NbBBLb    gagaaaaacccgaagggataccttgtcttcgatccttatggcggagccatggacaagatt  1275
NtBBLc    gagaaaaaccccaatgcattccttgtgttcgatccatatggcggagccatggacaaaatt  1263
NtBBLe    gagaaaaaccccaagtcattccttgtcttcgatccttatggcggagtcatggacaagatt  1275
NtBBLa    gagaaaaacccgaagggatatcttgtctttgatccttatggcggagccatggacaagatt  1251
NtBBLb    gagaaaaacccgaagggatatcttgtctttgatccatatggcggagccatggacaagatc  1278
NbBBLd    gagaaagaacccaatggacatgtcatatttgaccccttatggtgcagccatgcagagaatt  1275
NbBBLd'   gagaaagaacc-aatggacatgttatctttgaccccttatggtgcagccatgaagagaatt  1272
NtBBLd1   gagaaagaacccaacggacatgtcatcttggaccccttatggtggagccatgcaaagaatt  1275
NtBBLd2   gagaaagaaccaaatggacatgtcatctttgaccccttatggtgcagccatgcagagaatt  1275

Consensus agtgatcaagctattgctttccctcatagaaagggtaaccttttcgcrattcaatatcta  1330
NbBBLc    agtgatcaagctattcctttccctcatcgaaagagtaaccttttcgccgttcaatattat  1338
NbBBLa    agtgatcaagctattgctttccctcatagaaaaggtaaccttttcgcgattcagtatctg  1323
NbBBLb    agtgatcaagctattgctttccctcatagaaaaggtaaccttttcgcgattcagtatctg  1335
NtBBLc    agtgctcaagctattgctttccctcatcgaaagggtaaccttttcgcaattcaatattat  1323
NtBBLe    agtgatcaagctattgctttccctcatcgaaagggtaaccttttcgcggttcaatattat  1335
NtBBLa    agtgatcaagctattgctttccctcatagaaaaggtaaccttttcgcgattcagtatcta  1311
NtBBLb    gatgatcaagctattgcgttccctcatagaaagggtaatcttttcgcgattcaatatcta  1338
```

```
NbBBLd    agcgaggaagctattgctttccctcatagaaagggtaacctattcggaattcaatatcta  1335
NbBBLd'   ggcgaggaagctattgctttccctcatagaaagggtaatcttttcggaattcaatatcta  1332
NtBBLd1   agtgaggaagctattgctttccctcatagaaagggtaacctttcggaattcaatatcta  1335
NtBBLd2   agcgaggaagctattgctttccctcatagaaagggtaacctattcagaattcaatatcta  1335

Consensus gcastgtggaawgaagaggacgatwa---------caagagcaa---cgkgtacatagag  1378
NbBBLc    gcagtgtggaacgaagaggacgatgc---------------cga---cgagtacttagag  1380
NbBBLa    gcacagtggaatgaagaggacgatta---------catgagcga---cgtttacatggag  1371
NbBBLb    gcacagtggaatgaagaggacgatta---------catgagcga---cgtttacatggag  1383
NtBBLc    gcacagtggaacgaagaggacgatgc---------caagagcaa---cgagcacatagag  1371
NtBBLe    gcattttggaacgaagaggacgatgc---------caagagcaa---cgagtacatagag  1383
NtBBLa    gcacagtggaatgaagaggacgatta---------catgagcga---cgtttacatggag  1359
NtBBLb    gcacagtggaatgaagaggacgatta---------caagagcga---tgtttacatggag  1386
NbBBLd    gtagtgtggaaagaaaaggacaataataatattgccaagagcaa---tgggtacatagag  1392
NbBBLd'   gtagtgtggaaagaaaaggacaataataatattgccaagaggaa---tggatacatagag  1389
NtBBLd1   gtagtgtggaaagaaaaggacaataataatattgtcaagagcaatattgggtacatagag  1395
NtBBLd2   gtagtgtggaaagaaaaggacaataataatattgccaagagcaa---tgggtacatagag  1392

Consensus tggataagaggatttacaatacaatggcgccctttgtttcaagctcgccaaggggagct  1438
NbBBLc    tggataagaggattttataataaaatggcgcctttgtttcaagctcgccaaggggagct  1440
NbBBLa    tggataagagggttttacaatacaatgaccccatttgtgtcaagctcgccaaggggagct  1431
NbBBLb    tggataagaggattttacaatacaatgacgccatttgtttcaagctcgccaaggggagct  1443
NtBBLc    tggataagaggattttacaataaaatggcgcctttgtttcaagctcgccaaggggagct  1431
NtBBLe    tggacaaggggattttacaataaaatggcgcctttgtttcaagctcgccaaggggagct  1443
NtBBLa    tggataagaggattttacaatacaatgacgccctttgtttcaagctcgccaaggggagct  1419
NtBBLb    tggataagaggattttacaatacaatgacgccctttgtttcaagctcgccaaggggagct  1446
NbBBLd    tggataagagagttttacaatacaatggcaccccatgtttcaacttcacctagggcagct  1452
NbBBLd'   tggataagagagctttacaatacaatggcaccccatgtttcaagttcaccaagggcagct  1449
NtBBLd1   tggataagagagttttacaatacaatggcaccccatgtttcaagttcacctagggcagct  1455
NtBBLd2   tggataagagagttttacaatacaatggcaccccatgtttctagttcacctagggcagct  1452

Consensus tatatcaactacttggatatggatcttggagtgaatatggtcgacgactacttaytgcga  1498
NbBBLc    tatatcaactacttggatatggatcttggagtgaacatggaccatgactacttactgcga  1500
NbBBLa    tatatcaactacttggatatggatcttggagtgaatatggtcgacaactacttattgcta  1491
NbBBLb    tatatcaactacttggatatggatcttggagtgaatatggtcgacaactacttattacga  1503
NtBBLc    tatgtcaactacttggatatggatcttggaatgaatatggacgacgactacttactgcga  1491
NtBBLe    tatatcaactacttggatatggatcttggagtgaatatggacgacgactacttactgcga  1503
NtBBLa    tatatcaactacttggatatggatcttggagtgaatatggtcgacgactacttattgcga  1479
NtBBLb    tatatcaactacttggatatggatcttggagtgaatatggacgacgactacttactgcga  1506
NbBBLd    tatgtcaactatatggatcttgaccttggagtga------tggacggctacttaatgcta  1506
NbBBLd'   tatgtcaactacatggatctggaccttacaa-----------------------------  1480
NtBBLd1   tatgtcaactacatggatctggaccttggagtga------tggacgactacttattgcca  1509
NtBBLd2   tatgtcaactatatggatctggaccttggagtga------tggacgactacttaatgcta  1506

Consensus aatgctagtagn---nnttcttcttcctctgttgatgctgtggagagagctagagcgtgg  1555
NbBBLc    aatacta------------cttcttcctctgttgatgctgtggagagagctagagcgtgg  1548
NbBBLa    aatgctagcagt---agttcttcttcctctgctgatgctgtggaaagagctagagcatgg  1548
NbBBLb    aatgctactagt---agttcttcttcctctgttgatgctgtggagagagctagagcgtgg  1560
NtBBLc    aatgctagtagt---cgttattcttcctctgttgatgctgtggagagggctagagcatgg  1548
NtBBLe    aatgctagtagtcgtagttcttcttcctctgttgatgctgtggagagagctagagcgtgg  1563
NtBBLa    aatgctagtagcagtagtccttcttcctctgttgatgctgtggagagagctagagcgtgg  1539
NtBBLb    aatgctagtagtcgtaattcttcttcctctgttgatgctgtggagagagctagagcgtgg  1566
NbBBLd    aatactagtac------tactgcctcttctgatcatgccgtggagagagcaagggtctgg  1560
NbBBLd'   ------------------------------------------------------------  1480
NtBBLd1   tgtactagtac------tactgcgtctgctaatcatgccgtggagagagcaagggtctgg  1563
NtBBLd2   aatactagtat------tactgcctctgctgatcatgccgtggagagagcaagggtctgg  1560

Consensus ggtgaaawgtatttcttgmataactatgataggttggttaaagctaagacacaaattgat  1615
NbBBLc    ggtgaaaagtatttcttgaacaactatgataggttggttaaagctaagacacaaattgat  1608
NbBBLa    ggtgaaatgtatttcttgcataactatgataggttggttaaagctaagacacaaattgat  1608
NbBBLb    ggtgaaatgtatttcttgcataactatgataggttggttaaagctaagacacaaattgat  1620
NtBBLc    ggtgaaaagtatttcttgaataactatgataggttggttaaagctaagacaaaaattgat  1608
NtBBLe    ggtgaaatgtatttcttgcataactatgataggttggttaaagctaagacacaaattgat  1623
NtBBLa    ggtgagatgtatttcttgcataactatgataggttggttaaagctaagacacaaattgat  1599
NtBBLb    ggtgaaatgtatttcttgcataactatgataggttggttaaagctaagacacaaattgat  1626
NbBBLd    ggtgaaaagtatttcttgaataactatgatagattggtcaaagctaagacaaaaattgac  1620
NbBBLd'   ------------------------------------------------------------  1480
NtBBLd1   ggtgaaaagtatttcttgaataactatgatagattggtcaaagctaagacaaaaattgac  1623
NtBBLd2   ggtgaaaagtatttcttgaataactatgatagattggtcaaagctaagacaaaaattgac  1620

Consensus ccactaaatgtttttcgacatgaacagagtattcctcctatgcttggttcaacgcaagag  1675
NbBBLc    ccactaaatgtttttcgacatgaacagagtattcctcctatgcttggttcaacgcaagac  1668
NbBBLa    ccactaaatgtttttcgacatgaacagagtattcctcctatgcgtggttcaacgcaagag  1668
NbBBLb    ccactaaatgttttccgccatgaacagagtattcctcctatgcttggttcaacgcaacag  1680
NtBBLc    ccactaaatgtttttcgacatgaacagagtatacctcctacgcttggttcaacgcaagag  1668
NtBBLe    ccactaaatgtttttcgacatgaacagagtatacctcctatgcttggttcaacgcaagag  1683
NtBBLa    ccactaaatgtttttcgacatgaacagagtattcctcctatgcttggttcaacgcaagag  1659
NtBBLb    ccacttaatgtttttcgacatgaacagagtatacctcctatgcttggttcaacgcaagag  1686
NbBBLd    ccactaaacgtttttcgacatcaacagggcatccctcctattttcgcctcaatgcaagag  1680
NbBBLd'   ------------------------------------------------------------  1480
```

```
NtBBLd1   ccactaaacgtttttcgacatcaacagggcatccctcctttgttcgcctcaatgcaagag  1683
NtBBLd2   ccactaaacgtttttcgacatcaacagggcatccctcctatgttcgcctcaatgccagag  1680

Consensus cataagtatagtagtgaatga                                         1696
NbBBLc    aataagtatagcagtgaatga                                         1689
NbBBLa    aacaagtatagtagcgaatga                                         1689
NbBBLb    cataagtatagtagtgaatga                                         1701
NtBBLc    cataactacagtagtgaatga                                         1689
NtBBLe    ca------cagtagtgaatga                                         1698
NtBBLa    cacaagtatagcagtgaatga                                         1680
NtBBLb    cataagtacagtagtgaatga                                         1707
NbBBLd    catacctacagtagtaaatga                                         1701
NbBBLd'   -------------ctacttaa                                         1488
NtBBLd1   tatacctatagtagtaaatga                                         1704
NtBBLd2   catacctatagtagtaaatga                                         1701
```

**Figure S3. Analysis of (*S*)- and (*R*)-nicotine in leaves of the quintuple *NbBBL* mutant (line 102) in comparison to control lines (WT and Cas9).** Traces correspond to extracted ion chromatograms (nicotine, $[M+H]^+$) resulting from chiral LC-MS analyses (Lux® 3 µm AMP column). Traces on the upper half of the figure are from uninduced plants, while traces on the lower half are from the same plants 5 days after induction with MeJa. Higher injection volumes were used for all uninduced samples (10 µL compared to 2 µL) to obtain comparable peak sizes. A total of four to five biological replicates were analyzed. Traces on the left column show the results of running a racemic nicotine standard and a (*S*)-nicotine standard along with the uninduced and induced samples.

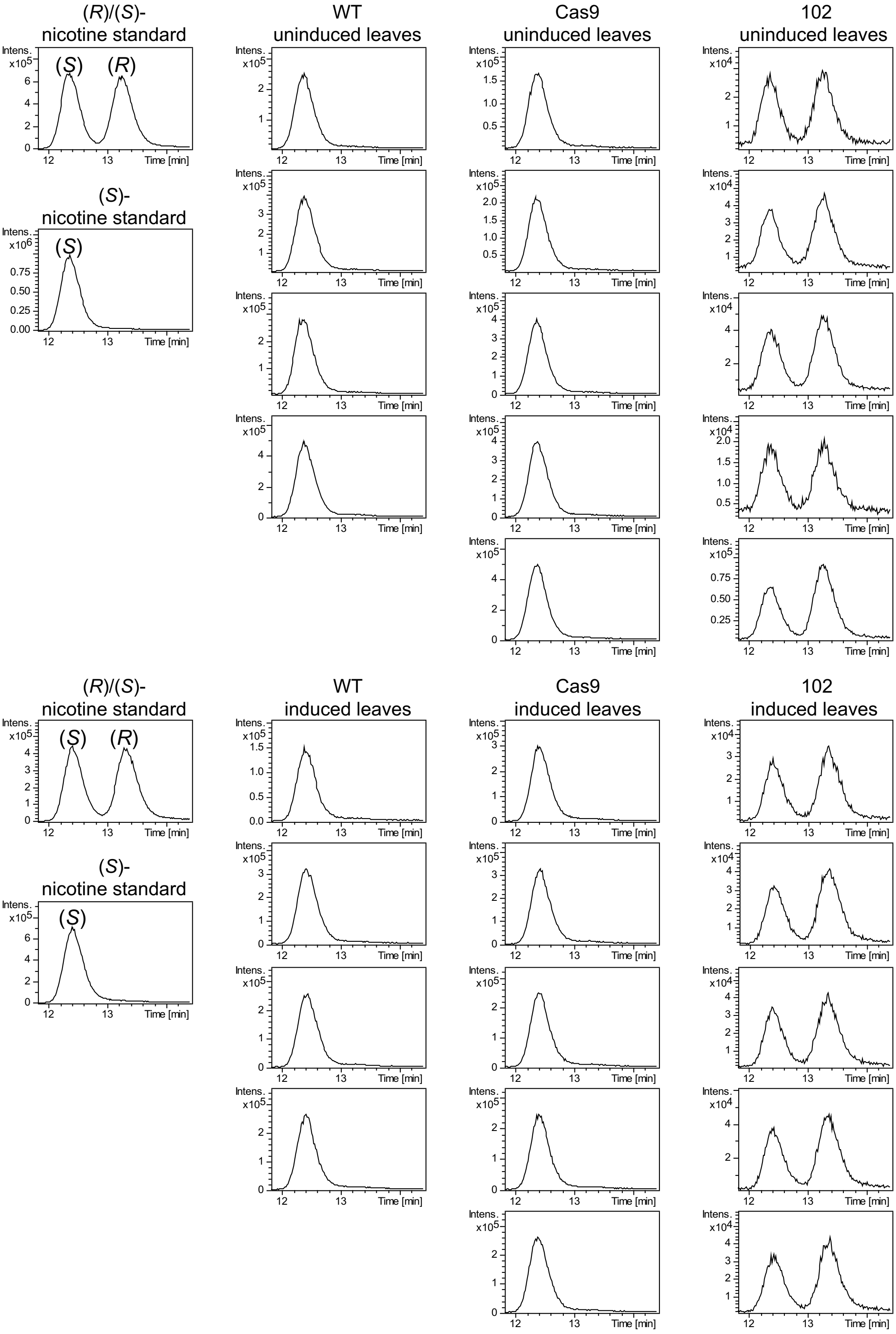

**Figure S4. DMN accumulation in roots of the quintuple *NbBBL* mutant (line 102) in comparison to two control lines (WT and Cas9), as analyzed by LC-MS.** Seedlings were grown under hydroponic conditions, and roots were harvested 5 days after induction with MeJa. Traces are extracted ion chromatograms ([M+H]$^+$) corresponding to nicotine (black) or DMN (blue). A total of five to six biological replicates per line were analyzed. The two traces at the top correspond to a (*S*)-nicotine standard and a DMN standard.

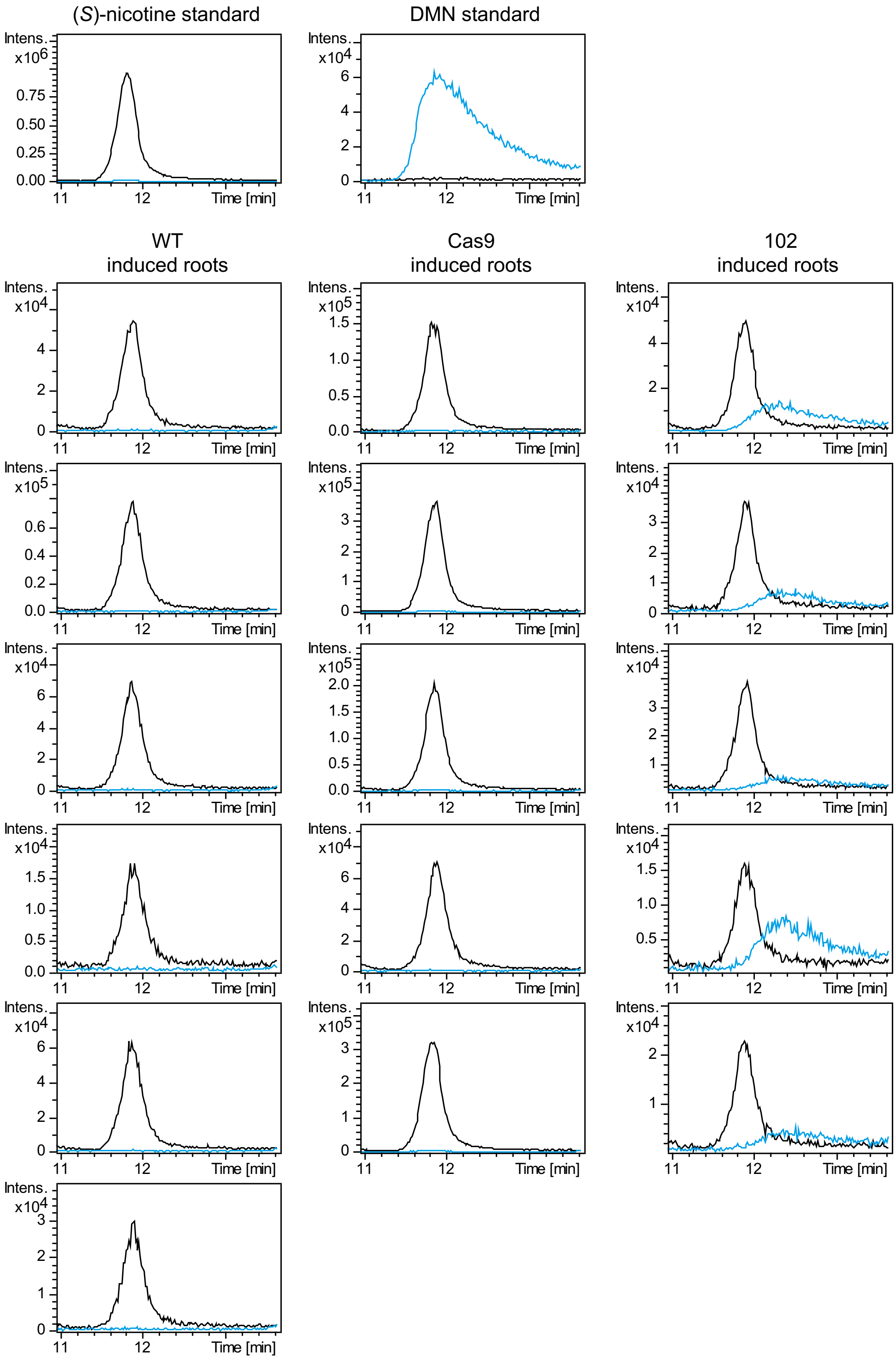
